# Multiplexed *in situ* protein imaging using DNA-barcoded antibodies with extended hybridization chain reactions

**DOI:** 10.1101/274456

**Authors:** Yu Wang, Yitian Zeng, Sinem K. Saka, Wenxin Xie, Isabel Goldaracena, Richie E. Kohman, Peng Yin, George M. Church

**Affiliations:** Wyss Institute for Biologically Inspired Engineering, Harvard University, Boston, USA; Department of Genetics, Harvard Medical School, Boston, USA; Department of System Biology, Harvard Medical School, Boston, USA

## Abstract

Immunofluorescence (IF) imaging using antibodies to visualize specific biomolecules is a widely used technique in both biological and clinical laboratories. Standard IF imaging methods using primary antibodies followed by secondary antibodies have low multiplexing capability due to limited availability of primary antibodies raised in different animal species. Here, we used a DNA-based signal amplification method, Hybridization Chain Reaction (HCR), to replace secondary antibodies to achieve multiplexed imaging using primary antibodies of the same species with superior signal intensity. To enable imaging with DNA-conjugated antibodies, we developed a new antibody staining protocol to minimize nonspecific binding of antibodies caused by conjugated DNA oligonucleotides. We also expanded the HCR hairpin pool from previously published 5 to 13 for highly multiplexed *in situ* imaging. We finally demonstrated multiplexed *in situ* protein imaging using the technique in both cultured cells and mouse retina sections.

## Introduction

Immunofluorescence imaging with antibodies has become a standard tool used to localize proteins *in situ*. A typical IF protocol involves labeling specific targets with primary antibodies, followed by signal amplification using fluorophore-conjugated secondary antibodies targeting the primary antibodies. The secondary antibodies are raised against the constant regions of primary antibodies, and are species-specific (e.g. anti-mouse secondary antibodies). This poses a significant challenge for multiplexed imaging, as it necessitates that primary antibodies from different species or different subclasses (in the case of mouse antibodies) are used to multiplex. Unfortunately, most of the validated antibodies (particularly monoclonal antibodies) are produced either in mice or rabbits, hindering flexible antibody selection. Primary antibodies directly conjugated with fluorophores have been used to bypass this issue; however, a lack of signal amplification from secondary antibodies typically restricts this method to visualizing only high abundance targets. While sequential immunostaining and antibody removal can also be used to bypass the antibody species limitation^1-4^, it is time-consuming and could deteriorate the sample integrity over multiple rounds of staining.

In order to address the aforementioned challenges, we integrated a DNA-based signal amplification method, Hybridization Chain Reaction (HCR)^5-8^, with IF to achieve multiplexed imaging with primary antibodies of the same species. HCR uses a single DNA initiator sequence to trigger the assembly of a linear DNA structure by iterative HCR hairpin openings (**Supplementary Fig. 1**)^5-8^. Each hairpin is labeled with a fluorophore, and the signal is amplified by hairpin stacking. For protein target imaging, primary antibodies are conjugated with distinct DNA initiator sequences and are applied together to label multiple targets in the sample. HCR hairpins labeled with spectrally distinct fluorophores are then used to simultaneously amplify signals for all targets, followed by fluorescence microscopy imaging (**Fig. 1**).

**Figure 1.**
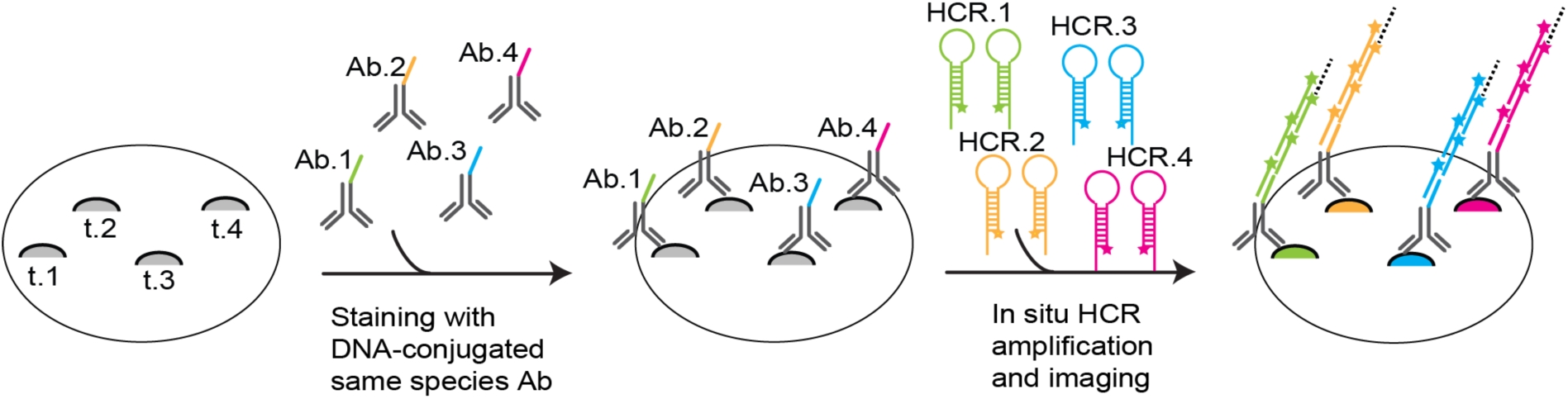
Schematic of imaging using DNA-barcoded antibodies with extended HCR for multiplexing. Targets (t.1-t.4) within biological samples are labeled with antibodies that are conjugated with distinct HCR DNA initiators (Ab.1-Ab.4). The signals are amplified simultaneously through orthogonal HCR reactions (HCR.1-HCR4), followed by spectrally multiplexed fluorescence microscopy imaging.

HCR has been previously used to detect small molecules (e.g. ATP) and RNA^2,4^. The method was more recently used to amplify protein signals with DNA-conjugated secondary antibodies^9-11^. When we tried to apply this approach for DNA-conjugated primary antibodies to enable higher multiplexing, we observed strong nonspecific fluorescence signals, particularly in the cell nuclei. Here, we developed a new antibody staining protocol to minimize nonspecific binding of DNA-conjugated antibodies by blocking the hybridization of conjugated DNA to endogenous DNA/RNA molecules and by reducing the electrostatic interactions between conjugated negatively charged DNA to endogenous positively charged molecules. To enable highly multiplexed detection, we designed and screened 15 HCR pairs in addition to the 5 commercially available pairs, and successfully expanded the pool of validated HCR pairs. We finally demonstrated multiplexed *in situ* imaging using DNA-barcoded antibodies with HCR in various sample types.

## Results

### Reduction of nonspecific DNA-conjugated antibody staining

We first directly conjugated HCR-initiator DNA sequences to primary antibodies through covalent chemical modification^12^. We tested the labeling specificity of DNA-modified primary antibodies by staining the antibodies in cultured BS-C-1 cells followed by HCR amplification. Surprisingly, we observed strong nonspecific signals, especially inside the nuclei (**Fig. 2a**). The nonspecific signal persisted when DNA-conjugated antibodies were indirectly detected by conventional fluorophore-conjugated secondary antibodies instead of the hairpins, but was absent in the negative samples where HCR hairpins were applied without the antibodies, suggesting the signals were caused by nonspecific binding of DNA-conjugated antibodies.

**Figure 2.**
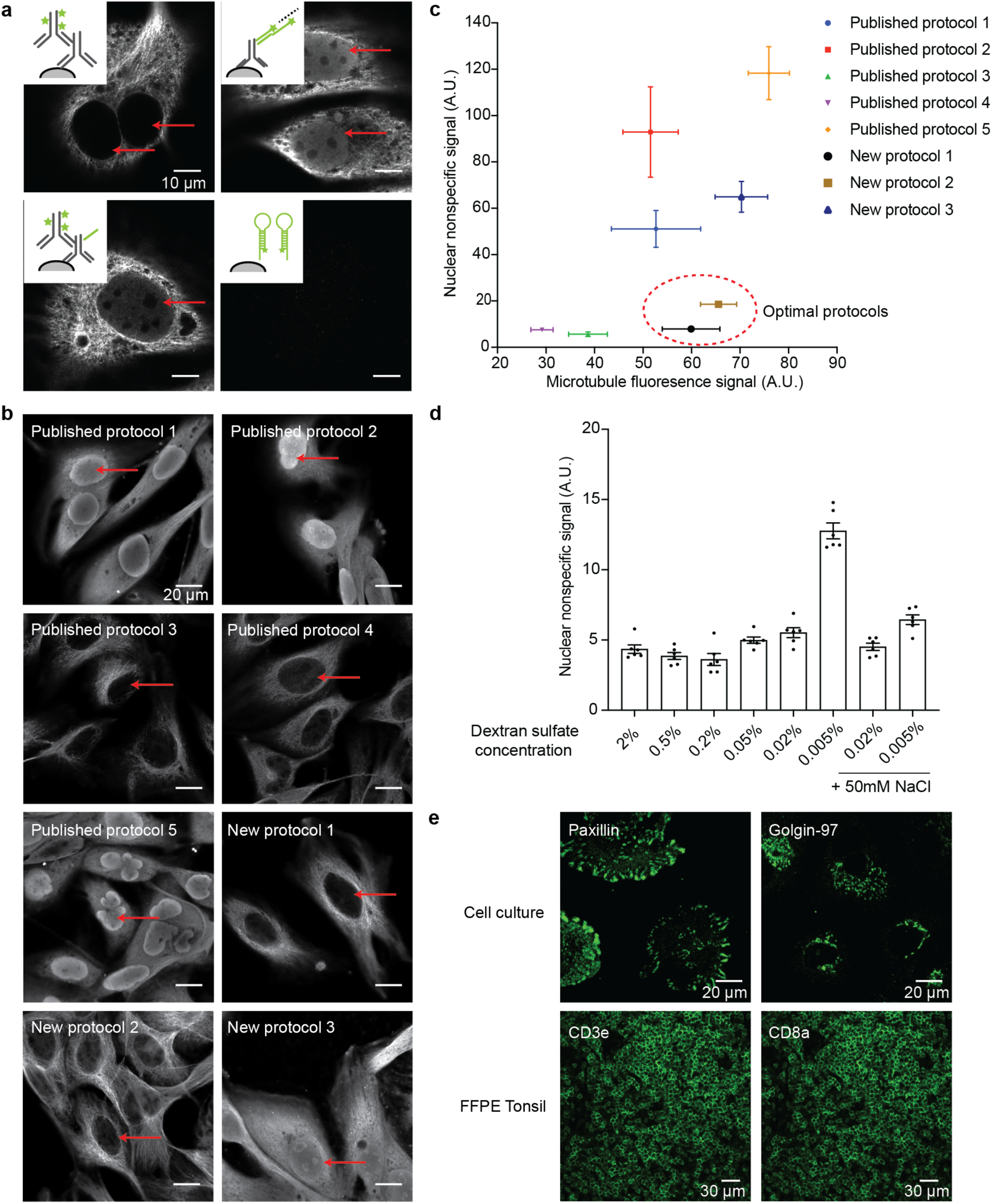
Development of DNA-conjugated antibody staining protocols with minimized nonspecific signals. **a)** Nonspecific nuclear signals came from the nonspecific binding of DNA-conjugated antibodies in the nuclei. BS-C-1 cells were fixed and stained with regular or HCR-B1I1-conjugated anti-β-Tubulin E7 antibodies to visualize cytoplasmic microtubules. The schematics for each experiment were shown in the top-left corner of each image. **b)** Comparison of different antibody staining methods for nonspecific nuclear signal reduction efficiency. BS-C-1 cells were stained with HCR-B1I1-conjugated anti-β-Tubulin E7 antibodies with protocols described in Supplementary Table 1. Representative cell nuclei were indicated with red arrows. **c)** Nonspecific nuclear signals and specific microtubule signals were quantified and plotted in a 2-D plot. The optimal protocols had minimal nonspecific nuclear signals but retained strong microtubule signals. Error bar is SEM and n = 6 areas. **d)** Titration of dextran sulfate in the antibody incubation buffer. **e)** Demonstration of the finalized staining protocol with DNA-conjugated Paxillin/Golgin-97 antibodies in BS-C-1 cells and CD3e/CD8a antibodies in FFPE human tonsil sections.

We decided to optimize the antibody staining protocol to reduce the nonspecific binding of DNA-conjugated antibodies. We curated a list of staining protocols that have been used in previous literature involving DNA-conjugated antibodies, and developed three new protocols by analyzing the components in the published protocols (**Supplementary Table 1**). We hypothesized that the nonspecific binding of DNA-conjugated antibodies was mainly caused by 1) the hybridization of conjugated DNA oligonucleotides with intracellular DNA/RNA molecules and 2) the electrostatic interactions between negatively charged DNA oligonucleotides with cellular positively charged molecules such as histone proteins. In the new protocols, we tested the following strategies to address these two causes: 1) convert the single-stranded conjugated DNA to double-stranded by hybridization of complementary DNA before antibody incubation to prevent the binding of DNA-conjugated antibodies to intracellular nucleic acids. 2) add negatively charged polymer dextran sulfate to compete for electrostatic interactions with positively charged molecules. 3) add sheared salmon sperm DNA to compete with conjugated DNA on antibodies. 4) increase the ionic strength of the buffer by adding Na^+^ to neutralize the negative charge of DNA. Bovine serum albumin (BSA) or normal mouse/rabbit IgG and detergent Triton X-100 were added in all solutions to block nonspecific interactions as standard immunostaining protocols.

We conjugated two antibodies (anti-α-Tubulin YL1/2 clone and anti-β-Tubulin E7 clone) with HCR initiators, and compared the intensity of nonspecific nuclear signals as well as microtubule fluorescence signals in different antibody staining protocols (**Fig. 2b, c and Supplementary Fig. 2a-c**). We noticed the degree of nonspecific signals was antibody-dependent, and anti-β-Tubulin E7 antibody showed more severe nonspecific signals than anti-α-Tubulin YL1/2 antibody. In the case of anti-β-Tubulin E7 antibody, Published protocol 1, 2, 5 and New protocol 3 (see **Supplementary Table 1**) failed to reduce nonspecific nuclear signals, whereas Published protocol 3 and 4 eliminated nonspecific nuclear signals but also reduced microtubule signals. New protocol 1 and 2 yielded more optimal staining with minimized nuclear signals and retained microtubule signals (**Fig. 2b, c and Supplementary Fig. 2a**). Similar results were observed for the anti-α-Tubulin YL1/2 antibody (**Supplementary Fig. 2b and c**). These two protocols shared the same components of complementary DNA oligonucleotides (blocks the DNA barcodes on antibodies) and dextran sulfate (masks electrostatic interactions with positively charged molecules) (**Supplementary Table 1**). We later showed that in the presence of complementary DNA oligonucleotides, sheared salmon sperm DNA was optional in the staining buffers (**Supplementary Fig. 2d**). The reduction of microtubule signals in Published protocol 3 and 4 was likely due to the high concentration of dextran sulfate, as dextran sulfate had been shown to alter antibody affinity^13^. Therefore, we titrated the dextran sulfate concentration in the antibody incubation buffer from 2% to 0.005% (w/v), and found that dextran sulfate was effective at blocking at concentrations as low as 0.02% (**Fig. 2d and Supplementary Fig. 2e**). Increasing the ionic strength of the buffer helped reduce the nonspecific nuclear signals at low dextran sulfate concentration (**Fig. 2d**). Dextran sulfate of different polymer size were commercially available, so we tested the efficacy of dextran sulfate of three different molecular weight (MW > 500 KDa, 9∼20 KDa and 6.5∼10 KDa). It could be seen that dextran sulfate of high MW (>500 KDa and 9∼20 KDa) had more consistent performance than the low MW version (**Supplementary Fig. 2f**).

Hence, our finalized buffer compositions are as follows. Blocking buffer: 1∼3% BSA + 0.1 mg/ml normal mouse/rabbit IgG + 0.1% Triton in 1× PBS (optional: 1 µM 50 nucleotide long poly TTG DNA sequences or 0.2 mg/ml sheared salmon sperm DNA); Antibody incubation buffer: 1∼3% BSA + 0.1 mg/ml normal mouse/rabbit IgG + 0.1% Triton + complementary DNA sequences with 1 µM for each sequence + 150 mM NaCl + 0.02%∼0.1% Dextran sulfate + 5 mM EDTA in 1× PBS. We demonstrated the finalized protocol with a few other antibodies in different sample types, including cell cultures and FFPE samples (**Fig. 2e**).

### Design and validation of new orthogonal HCR pairs

HCR-based signal amplification has the potential for highly multiplexed imaging by designing a pool of orthogonal HCR pairs, with each target amplified by one HCR pair. Five HCR pairs have been previously published and commercially available^7,8^. For higher multiplexing capability and more flexible HCR pair selection, we designed fifteen additional pairs of HCR hairpins and tested their performance and pairwise crosstalk *in silico* using NUPACK^14^. We then screened the leakage (i.e. hairpin assembly without initiators), amplification efficiency (i.e. hairpin assembly with cognate initiators), and pairwise crosstalk (i.e. hairpin assembly with noncognate initiators) *in vitro* for the new HCR hairpins using a gel shift assay (**Fig. 3a-d and Supplementary Fig. 3 and 4**). We selected eight pairs (B8, B9, B10, B11, B13, B14, B15 and B17) that had the best performance out of the fifteen new pairs of HCR oligonucleotides. B6, B12 and B18-20 had higher leakage than others *in vitro*, and were excluded for crosstalk analysis. We found that B8 initiator B8I2 had crosstalk with B7 hairpins, and B1 initiator B1I1 had crosstalk with B1hairpins. Therefore, we removed B7 and B16 from the HCR pair list.

**Figure 3.**
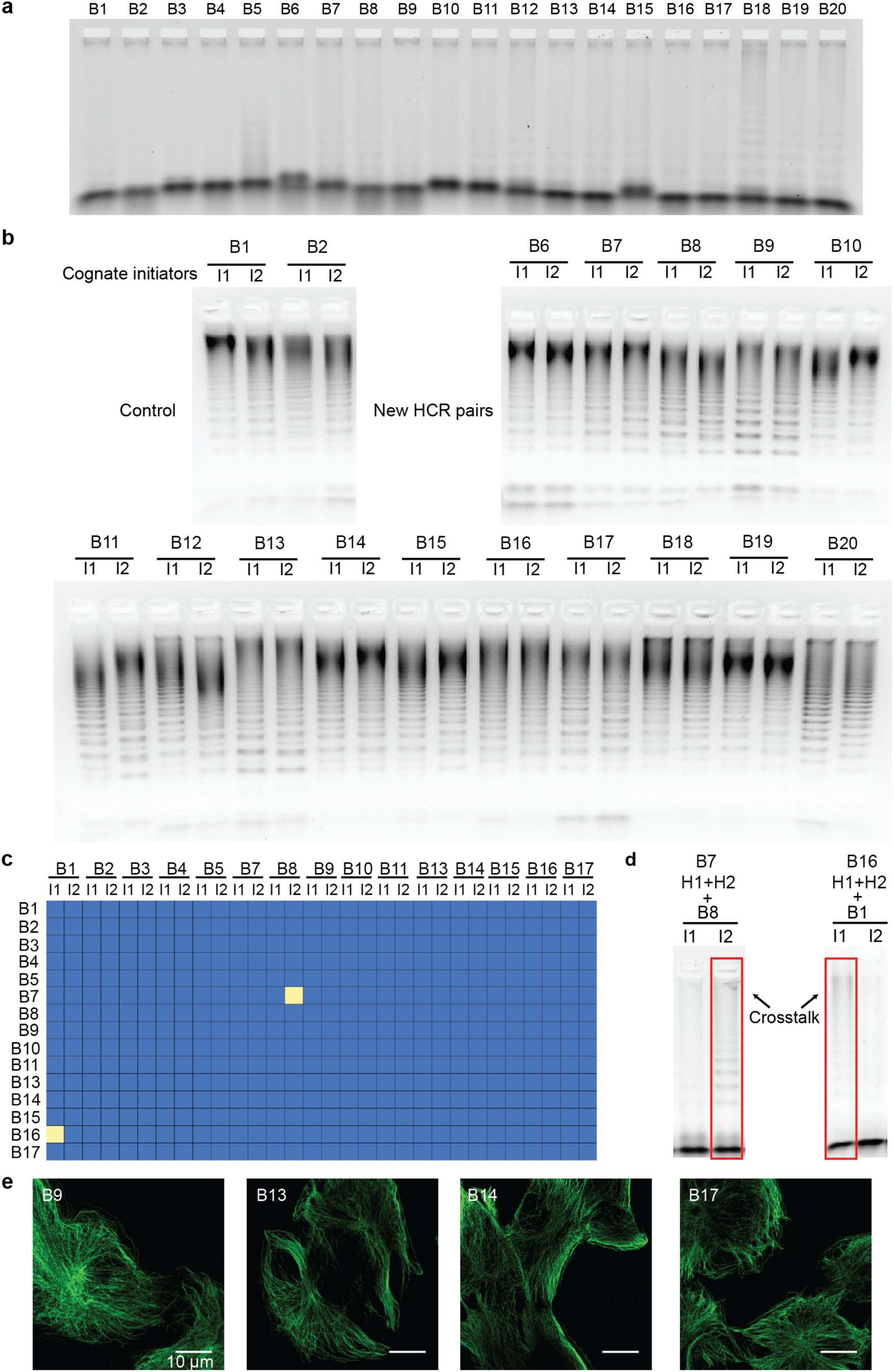
Validation of newly designed orthogonal HCR pairs. **a)** *In vitro* leakage analysis of newly designed HCR pairs. For each pair, 60 nM hairpins (H1 + H2) were added in a PCR tube without HCR initiator and left at room temperature for 24 hours. The results were visualized by running the products on agarose gels. **b)** *In vitro* amplification efficiency analysis of newly designed additional HCR pairs. For each pair, 500 nM of each HCR hairpin pair (H1 + H2) were added in a PCR tube with 25 nM of one of the two cognate initiators (I1 or I2). I1 refers to the initiator sequence that opens H2, and I2 refers to the initiator sequence that opens H1. The samples were left at room temperature for 24 hours. **c)** Summary of *in vitro* pairwise crosstalk analysis of 15 HCR pairs (excluding B6, B12, B18-20). For each reaction, 500 nM of each of the HCR hairpins (H1 and H2) was added in a PCR tube with 50 nM of the indicated initiator sequence, and left at room temperature for 24 hours to react. Reactions showing crosstalk were marked with yellow squares. The complete gel data are in the Supplementary Figure 4. **d)** The gel lanes with reactions showing crosstalk are marked with red boxes. **e)** *In situ* imaging of microtubules using newly designed orthogonal HCR pairs. Anti-β-Tubulin E7 antibodies were conjugated with newly designed HCR initiators (B9I1, B13I1, B14I1 and B17I1), and used to stain fixed BS-C-1 cells to visualize microtubule structures.

**Figure 4.**
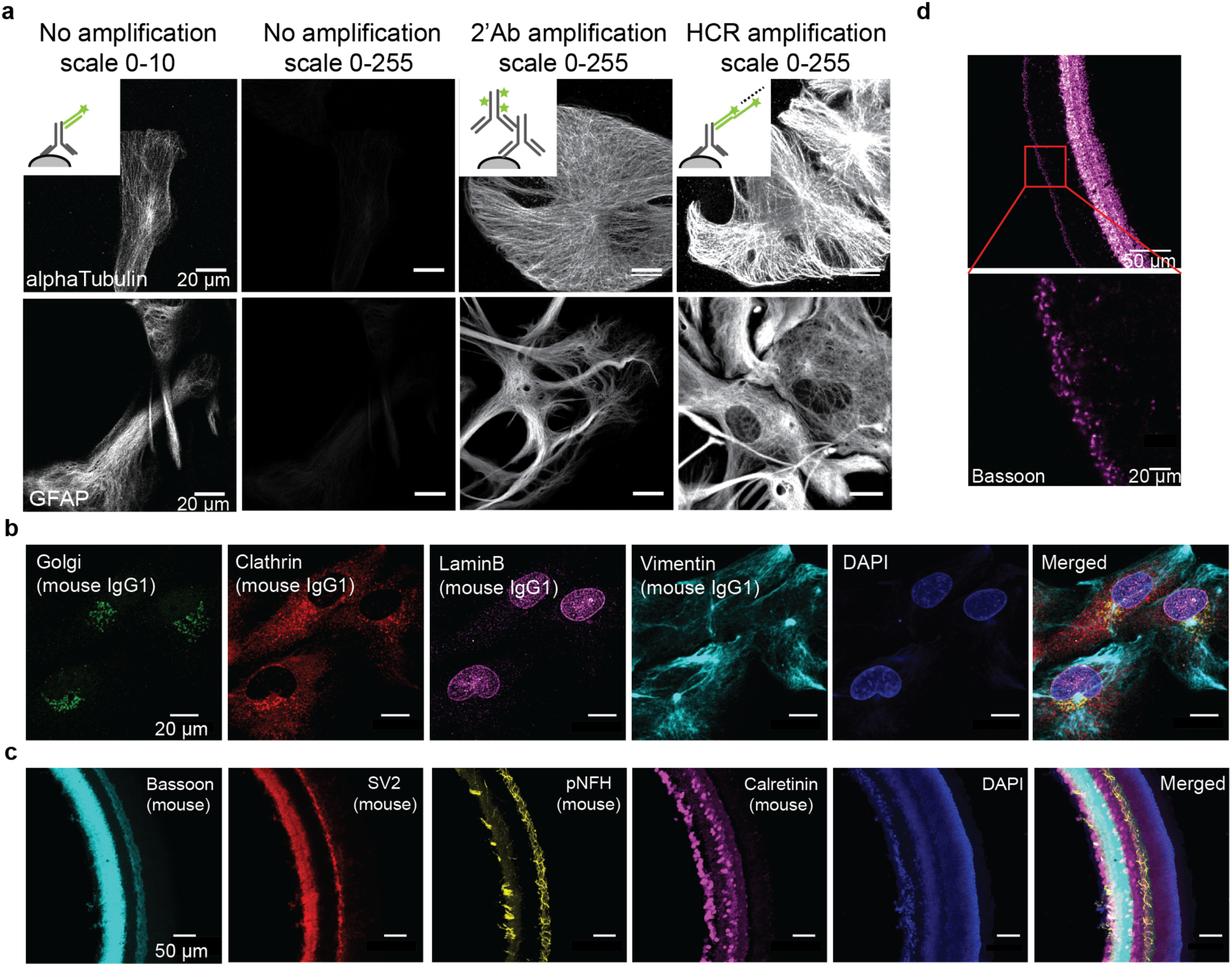
Multiplexed *in situ* protein imaging using HCR. **a)** Comparison of signals using conventional fluorophore-conjugated antibodies and HCR-based signal amplification. The schematics of each experiment were shown in the top-left corner of the images. In the upper panel, BS-C-1 cells were fixed and stained with HCR-B1I1-conjugated anti-α-Tubulin YL1/2 antibodies. In no amplification samples, the images were acquired using Alexa647-conjugated DNA strands that were complementary to the HCR initiator DNA sequences. Either Alexa647-conjugated secondary antibodies (2’Ab) or HCR hairpins were used to amplify the signals. Lower panel: primary mouse hippocampal neuron cultures were fixed and stained with HCR-B5I1-conjugated antibodies against GFAP. **b)** Multiplexed imaging using antibodies of the same species with HCR-based signal amplification in cultured BS-C-1 cells. Primary antibodies of mouse IgG1 subclass primary antibodies were conjugated with HCR-initiators (Golgi-B3I1, Clathrin-B2I1, LaminB-B4I1 and Vimentin-B5I1) and used to stain fixed BS-C-1 cells. DAPI was used to visualize the nuclei. **c)** Multiplexed imaging with the same species antibodies with HCR-based signal amplification in mouse retina sections. 40 μm mouse retina sections were stained with HCR initiator-conjugated primary antibodies against targets as labeled (Bassoon-B1I1, SV2-B2I1, pNFH-B4I1 and Calretinin-B5I1). **d)** Visualization of HCR amplified Bassoon signals in mouse retina sections. A 40 μm mouse retina section was stained with HCR B1-conjugated Bassoon primary antibodies followed by signal amplification using Alexa647-labeled HCR B1 hairpins.

We then validated four new HCR pairs (B9, B13, B14, B17) *in situ* by amplifying β-Tubulin signals in BS-C-1 cells (**Fig. 3e**). The results showed that all four of the new HCR pairs successfully assembled *in situ* to amplify signals from anti-β-Tubulin primary antibodies.

### Multiplexed *in situ* protein imaging using HCR

We then compared the signal level using HCR amplification with the unamplified case and commercial fluorophore-conjugated secondary antibodies. We stained cultured BS-C-1 cells with anti-α-Tubulin antibodies conjugated with HCR initiator B1I1- or B5I1, and primary neurons with anti-GFAP antibodies conjugated with HCR initiator B5I1 (**Fig. 4a**). The results showed that HCR outperformed commercial fluorophore-conjugated secondary antibodies and achieved better signal amplification. The quantification of signals was shown in Supplementary Figure 5. However, it should be noted that the degree of signal amplification using HCR depends on the concentration of HCR hairpins added in the solution. Higher concentration of HCR hairpins results in larger assembled DNA linear structures and hence stronger signals.

We then performed multiplexed imaging using the same species primary antibodies by conjugating distinct HCR initiator sequences to different primary antibodies. We first attached four mouse IgG (IgG1 subclass) primary antibodies targeting four cellular structure proteins (Golgin-97, Lamin B, Vimentin and Clathrin) with HCR initiators (B2I1 to B5I1). We then stained cultured BS-C-1 cells with the four antibodies, followed by HCR signal amplification and confocal microscopy imaging. We were able to get high-quality images for all four targets (**Fig. 4b**). We also performed multiplexed imaging in mouse retina tissue samples using four mouse IgG primary antibodies targeting Bassoon, SV2, pNFH (phosphorylated neurofilament heavy) and Calretinin (**Fig. 4c**). Bassoon and SV2 are two synaptic markers, and pNFH and Calretinin are two neuronal markers. Bassoon is located in the active zone of synapses, and is in low abundance^15^. The results showed that after HCR signal amplification, the Bassoon signals were clearly visible in 40 μm thick mouse retina sections (**Fig. 4d**).

### Discussion

Using antibodies to visualize specific targets *in situ* dates back to the 1940s when Dr. Albert Coons used fluorescein isocyanate-conjugated anti-pneumococcal serum to detect *Streptococcus pneumoniae* in formalin-fixed infected mouse tissues^16,17^. The technique has since been advanced with numerous developments, such as monoclonal antibodies and bright fluorophores. More recently, DNA-barcoded antibodies, which has significantly increased the multiplexing capability of immunofluorescence (IF) imaging methods, were introduced into the field, by us and several other groups ^10-12,15,18-24^. In addition to multiplexing, DNA-conjugated antibodies allow integrating DNA-base nanostructure assembly *for in situ* signal amplification.

The primary concern about conjugating DNA to antibodies is that the attached DNA can affect antibody binding affinity and specificity. Indeed, we observed strong nonspecific signals, particularly in the nuclei, with DNA-conjugated antibodies. The level of nonspecific signals was antibody-dependent with some antibodies showing higher nonspecific signals than the others. In addition, longer DNA sequences (e.g. ∼36-42 nucleotides in this paper) showed more severe nonspecific signals than shorter sequences (e.g. ∼10-11 nucleotides in DNA-Exchange-Imaging^20^). This could be the reason that the published protocols we tested failed to reduce nonspecific signals in our experiment. We reasoned that this nonspecific signal was caused by the hybridization of single-stranded DNA to intracellular nucleic acids and by the electrostatic interactions between negatively charged DNA molecules with positively charged cellular molecules. Therefore, we developed a new antibody staining protocol by introducing complementary DNA strands to block the nonspecific hybridization and negatively charged polymer dextran sulfate to compete for electrostatic interactions. We also optimized the protocol by tuning the concentration of sheared salmon sperm DNA, dextran sulfate and sodium chloride. The protocol worked reliably for the majority of antibodies we have tested (>80%), but we did observe some antibodies that failed to yield specific staining. For example, anti-TOM20 2F8.1 clone lost its binding affinity in the presence of dextran sulfate. We tested dermastan sulfate and polyacrylic acid as dextran sulfate alternatives but both polymers failed to effectively reduce nonspecific nuclear signals (unpublished observation). We also tested uncharged peptide nucleic acid (PNA) to replace DNA as docking strands on antibodies so that dextran sulfate could be eliminated from the protocol. Although this approach also showed promise in our preliminary studies, the high cost of PNA could expectedly hinder its wide adoption.

Another concern related to DNA-conjugated antibodies is the DNA to antibody ratio (i.e. the number of DNA molecules on each antibody). The likelihood of antibody specificity change is proportional to the number of DNA molecules on antibodies. In our method, we optimized the protocol to attach only 1-3 DNA oligonucleotides per antibody to avoid over-labeling, minimizing the chance of disrupting the paratopes. As a trade-off, the signal of DNA-conjugated antibodies decreases. Various DNA *in situ* amplification strategies can be used to enhance signals, such as HCR (Hybridization Chain Reaction), SABER (Signal Amplification by Exchange Reaction)^18^ and RCA (Rolling Circle Amplification)^27^. The difficulty of controlling enzymatic reactions *in situ* renders RCA incompatible with visualizing fine structures such as microtubules. HCR amplifies signals *in situ* by hairpin stacking, whereas SABER applies *in vitro* synthesized DNA concatemers. In HCR, the signal amplification fold can be tuned by adjusting the concentration of hairpins added in the reaction, whilst SABER can increase the signal amplification capability by iterative DNA branching. SABER has the advantage of being compatible with DNA-Exchange-Imaging and therefore is simpler for highly multiplexed imaging. However, the large size of DNA concatemers used in SABER may limit probe penetration into samples with densely packed environments, such as those in iterative expansion microscopy where double-layer polyacrylamide gels are formed^28^. In contrast, HCR employs small-size DNA hairpins, and hence more suitable for post-gel formation signal amplification in CLARITY^29^ and Expansion Microscopy^28,30-32^. HCR hairpin design and validation are more challenging, as it requires each hairpin pair efficiently amplify signals in the presence of cognate initiator sequences but also is sufficiently stable to have minimal leakage without cognate initiators. Indeed, we designed 15 additional hairpin pairs and only selected 8 out of the 15 to have a final list of 13 pairs. High throughput methods could be developed to assist screening to develop more HCR pairs.

The potential of DNA-conjugated antibodies goes beyond *in situ* imaging. Coupled with droplet- or microwell-based single-cell sequencing technique, DNA-conjugated antibodies have been used for highly multiplexed cell surface marker labeling^33^. With the increasing interest in using DNA-conjugated antibodies for biomedical research, we believe the methods described in this manuscript would not only provide a useful resource for the imaging field but also facilitate the development of future technologies.

## Methods

### Cultured cells preparation and staining

All animal procedures were in accordance with the National Institute for Laboratory Animal Research Guide for the Care and Use of Laboratory Animals and approved by the Harvard Medical School Committee on Animal Care and the Massachusetts Institute of Technology Committee on Animal Care.

Primary mouse hippocampal neuron cultures were prepared from postnatal day 0 or 1 mice and plated on eight-well ibidi glass-bottom chambers (ibidi, Cat. No. 80827) with a density of 10,000 - 15,000 cells per well. Cells were grown for 14 days before fixation. BS-C-1 cells were plated on eight-well ibidi glass-bottom chambers (10,000 cells per well) and grown for 24 hours. Cells were fixed using 4% paraformaldehyde (PFA) for 15 minutes at room temperature, followed by quenching in 50 mM NH4Cl or 1 TBS for 7 minutes.

The following is the finalized staining protocol. Cells were then permeabilized and blocked in 0.1% Triton X-100, 1-3% nuclease-free BSA (americanBIO, CAS 9048-46-8) and 0.1 mg/ml normal mouse/rabbit IgG for ∼1-2 hours. 0.2 mg/ml sheared salmon sperm DNA or 1 μM 50nt poly TTG sequences can be added but is optional. While the samples were in the blocking step, antibodies were incubated with complementary blocking DNA sequences individually (1 μL of 1 mM complementary DNA sequence was mixed with antibodies in 1 PBS; the amount of complementary DNA was calculated assuming the final antibody incubation reaction volume is 100∼125 μL.), and left at room temperature for 30 minutes. The complementary DNA sequences are 16 nucleotides long and each initiator sequence has two complementary DNA sequences that covers the middle 32 nucleotides of HCR initiator sequences. The antibodies were pooled together and added with 0.1% Triton X-100, 1-3% nuclease-free BSA, 0.1 mg/ml normal mouse/rabbit IgG, 0.02%-0.1% dextran sulfate (Millipore, S4030), 150 mM NaCl, 5 mM EDTA in 1× PBS. The sample was left at room temperature for 2 hours or overnight (add 0.05% sodium azide if left overnight). The sample was washed with washing buffer (0.1% Triton X-100, 1% nuclease-free BSA in 1× PBS) five times (brief wash for the first two washes and 10-30 minutes incubation for the other three washes). The samples were post-fixed using BS(PEG)_5_ (ThermoFisher, 21581) for 30 minutes to 1 hour to crosslink antibodies to the samples, followed by quenching in 1× TBS for 10 minutes. The hybridized complementary DNA sequences were removed by washing with 40% formamide in 0.1× PBS for 3 times with 10 minutes each. The sample was then washed with 5× SSC + 1% Tween 20 twice to remove formamide.

### Mouse retina section preparation and staining

Animals were given a lethal dose of sodium pentobarbital (120 mg/kg) (MWI, 710101) and enucleated immediately. Eyes were removed and fixed in PFA for 15-30 minutes. Following dissection, retinas were immersed in 30% sucrose overnight prior to freezing in TFM (EMS, 72592) and cryo-sectioned at 40 μm. Eight-well ibidi glass-bottom chambers were treated with poly-D-Lysine overnight, followed by PBS washes. Retina sections were attached in ibidi chambers, dried by plating on a heated block and stored at -20 °C. Sections were washed with 1× TBS + 0.3% Triton X-100 three times with 10 minutes per wash. They were then blocked and stained as above.

### Formalin-fixed paraffin-embedded (FFPE) tonsil sample preparation and staining

The specimens were obtained from the archives of the Pathology Department of Beth Israel Deaconess Medical Center under the discarded/excess tissue protocol as approved in Institutional Review Board (IRB) Protocol #2017P000585. 5 μm-thick sections were cut with a rotary microtome, collected in a water bath at 30°C, transferred to coated positively charged glass slides and baked at 60°C for 2 hours. Slides were then placed on a PT-Link instrument (Agilent) for deparaffinization, rehydration and epitope retrieval (with citrate buffer). Slides were held at 4°C in 1× PBS until staining.

For staining, sections were outlined with a hydrophobic pen (ImmEdge Hydrophobic Barrier PAP Pen, Vector Laboratories #H4000), and incubated in a humidified chamber. The samples were then blocked and stained as above.

### Antibodies

α-Tubulin (clone YL1/2, ThermoFisher MA1-80017), β-Tubulin (E7, produced in house, hybridoma from DSHB), GFAP (clone 2.2B10, ThermoFisher 13-0300), Golgi-97 (clone CDF4, ThermoFisher A-21270), Clathrin Heavy chain (clone X22, ThermoFisher MA1-065), Lamin B (clone L-5, ThermoFisher 33-2000), Vimentin (EncorBiotechnology MCA2A52), Bassoon (clone SAP7F407, Enzo ADI-VAM-PS003), SV2 (produced in house, hybridoma from DSHB), pNFH (EncorBiotechnology MCA-AH1), Calretinin (EncorBiotechnology MCA-6A9).

### Antibody-DNA conjugation

The conjugation involves crosslinking of thiol-modified DNA oligonucleotides to Lysine residues on antibodies. In brief, 250 μM 5’ thiol-modified DNA oligonucleotides (Integrated DNA Technologies) were activated by 100 mM DTT for 2 hours at room temperature in the dark and then purified using NAP5 columns (GE Healthcare Life Sciences, 17-0853-02) to remove excessive DTT. Antibodies formulated in PBS only were concentrated using 0.5 mL 50KDa Amicon Ultra Filters (EMDMillipore, UFC510096) to 2 mg/ml and reacted with maleimide-PEG2-succinimidyl ester crosslinkers (ThermoFisher 22102) for 2 hours at 4 °C. For every 100 μg antibody, 3.4 μl of 0.85 mg/ml DMF-diluted crosslinker was used. Antibodies were then purified using 0.5 mL 7kDA Zeba desalting columns (ThermoFisher, 89883) to remove excessive crosslinkers. Activated DNA oligonucleotides were incubated with antibodies (11:1 DNA: Antibody ratio) overnight at 4 °C. Final conjugated antibodies were washed using in 2 mL 50KDa Amicon Ultra Filters six times to remove non-reacted DNA oligonucleotides. Conjugated antibodies were kept at 4 °C.

### *In situ* HCR amplification

Fluorophore-conjugated HCR hairpins that were previously published were purchased from Molecular Instrument. Fluorophore (Alexa647)-conjugated newly designed B9, B13, B14 and B17 hairpins were purchased from IDT with dual HPLC purification. For antibody incubation condition testing and HCR signal amplification experiment, Alexa647-conjugated B1 or B5 hairpins were used. For multiplexing experiment, Alexa647-conjugated B1 hairpins, Alexa594-conjugated B2 hairpins, Alexa514-conjugated B3 hairpins, Alexa546-conjugated B4 hairpins, and Alexa488-conjugated B5 hairpins were used. Samples were blocked in amplification buffer (5× SSC buffer, 0.1% Tween 20 and 10% dextran sulfate) for 1 hour. Meanwhile, HCR hairpins were snap-cooled separately (heat hairpins at 95 °C for 90s in a PCR machine, immediately put hairpins on ice for 5 minutes, and then leave hairpins at room temperature for 30 minutes). Hairpins were then diluted in amplification buffer to 60 nM for each hairpin. Samples were incubated with HCR hairpins overnight at room temperature, and free hairpins were removed by three washes with 5 × SSCT (5 × SSC + 0.1% Tween 20). Hairpins sequences are listed in Supplementary Table 2.

### Confocal image acquisition

Samples were left in 5 × SSC buffer during image acquisition. All images were acquired using a Zeiss Axio Observer with LSM 710 scanning confocal system with either 20×/0.8 NA dry objectives or 63×/1.46 NA oil-immersion objectives. The images were 512×512 pixels and acquired at acquisition speed 7. Each image was acquired by averaging 2 images. In multiplexed imaging experiments, Alexa 488 was visualized using a 488 nm laser; Alexa 514 was visualized using a 514 nm laser; Alexa 546 was visualized using a 546 nm laser; Alexa 594 was visualized using a 594 nm laser; Alexa 647 was visualized using a 633 nm laser.

### Image analysis and quantification

All images were visualized and scaled using FIJI. For nonspecific nuclear signals, the fluorescence intensity of random regions of each nucleus was measured using FIJI. For signal amplification quantification, a binary mask was created for each image to represent the cellular structures using MATLAB, and the average fluorescence intensity (the sum of fluorescence signal within the binary mask / the total pixel number within the binary mask) was calculated. Background fluorescence intensity was calculated by averaging the fluorescence outside cells. The final fluorescence intensity is derived by average fluorescence intensity minus background fluorescence intensity. For microtubule fluorescence signal measurement, the background fluorescence intensity was not subtracted.

### HCR hairpin design

New HCR hairpins B6-B12 were designed using NUPACK multi-state design function and screened for crosstalk *in silico* using NUPACK analysis function. New HCR hairpins B13-B20 were designed by manually mutating B1-B5 sequences and screened for crosstalk *in silico* using NUPACK analysis function.

### Hairpin PAGE purification

For newly designed HCR hairpins used for *in vitro* analysis, standard desalting hairpin oligonucleotides (72 nt) were purchased from IDT. Denatured 7% PAGE gels were used for hairpin purification, in which 420 g/L urea was added as the denaturing reagent. 2× loading buffer (Biorad, 1610768) was added to hairpin samples, followed by sample denaturing with a thermocycler (heat at 85 °C for 4 minutes then quench on ice for 5 minutes). Samples were loaded into the gel and ran at 150 V for 20 minutes then at 200 V for 90 minutes in a 50 °C water bath system. After electrophoresis, hairpin gel bands were cut out of the gel while visualizing their location using a short wavelength UV lamp while placing the gel above silica gel. Product was extracted from the excised gels by smashing the gel completely in a 1.5 mL Eppendorf tube, adding 1× TE buffer to the smashed gel, freezing at -20 °C for 30 minutes, and then rotating at 4 °C overnight. The gel was then centrifuged at 18,000 g for 10 minutes. To increase the DNA recovery rate, the supernatant was collected and the pellet was washed once with water. The combined supernatant was combined with 0.1 volume of 3 M NaAc (pH=5.5) and 3 volumes of ethanol and then put at -80 °C for 30 minutes. The sample was centrifuged at 18,000 g for 10 minutes, and the supernatant was carefully discarded to collect the white pellet. The pellet was washed once with 75% ethanol and then with 100% ethanol without disturbing the pellet. The sample was left to air dry for 5 minutes before being finally dissolved in ultrapure water.

### HCR *in vitro* leakage test

PAGE purified hairpins were first annealed in advance (95 °C for 3minutes, then decrease 0.1 °C per second till 10 °C). The two hairpins (H1 and H2) were then snap-cooled separately in 5× SSC (Heat hairpins at 95 °C for 90 seconds, and immediately put on ice for 5 minutes. The hairpins were then left at room temperature for 30 min). Hairpin were then mixed together and diluted with 5× SSCT (5× SSC + 0.1% Tween 20) to their final concentration. The samples were left at room temperature for 24 hours and run in a 2% agarose gel containing 1/10000 SYBR Gold (ThermoFisher S11494) at 90V for 30 min, then 120V for 90 minutes. The products were visualized using a gel scanner.

### HCR *in vitro* amplification test

HCR hairpins were snap-cooled separately in 5× SSC, mixed together, and diluted with 5× SSCT to a final concentration of 500 nM for each hairpin. Initiators were added to each reaction to a final concentration of 25 nM. The samples were left at room temperature for 24 hours and then ran in a 2% agarose gel (with 1/10000 SYBR Gold staining) at 90V for 30 minutes and then at 120V for 90 minutes. The products were visualized using a gel scanner.

### HCR *in vitro* crosstalk test

HCR hairpins were snap-cooled separately in 5× SSC, mixed together, and diluted with 5 SSCT to a final concentration of 500 nM for each hairpin. The initiator was added in the reaction at a final concentration of 50 nM. The samples were left at room temperature for 24 hours and then ran in a 2% agarose gel (with 1/10000 SYBR Gold staining) at 90V for 30 minutes and then at 120V for 90 minutes. The products were visualized using a gel scanner.

## Acknowledgements

We thank Dr. Demian Park for providing primary mouse hippocampal neuron cultures. We thank Dr. Sylvain Lapan for providing mouse retina samples. We thank Dr. German Pihan for providing FFPE samples. This work is supported by NIH grants (RM1HG008525, R01NS083898, R01MH113279) and IARPA grant (IARPA MICrONS; D16PC0008) to G.M.C and NIH grants (1RO1EB018659 and 1UG3HL145600/HuBMAP) to P.Y.. W.X. is partially supported by Fudan University visiting undergraduate student funding; Y.Z. is partially supported by Tsinghua University visiting undergraduate student funding.

## Author contributions

Y.W. conceived the study, designed and performed the experiments, analyzed the data. S.K.S contributed to the development of antibody staining protocols. W.X., Y.Z. and I.G. performed the experiments. R.E.K. and P.Y. supervised the project. G.M.C. conceived the study and supervised the project. Y.W. and G.M.C. wrote the paper with input from all authors. All authors reviewed and edited the paper.

## Competing interest

G.M.C. is a cofounder of Readcoor, Inc. and P.Y. is a cofounder of Utivue, Inc.

## Reference

1 Pirici, D. et al. Antibody elution method for multiple immunohistochemistry on primary antibodies raised in the same species and of the same subtype. J Histochem Cytochem 57, 567–575, doi: 10.1369/jhc.2009.953240 (2009).

2 Gerdes, M. J. et al. Highly multiplexed single-cell analysis of formalin-fixed, paraffin-embedded cancer tissue. Proc Natl Acad Sci U S A 110, 11982–11987, doi: 10.1073/pnas.1300136110 (2013).

3 Lin, J.-R. et al. Highly multiplexed immunofluorescence imaging of human tissues and tumors using t-CyCIF and conventional optical microscopes. eLife 7, doi: 10.7554/eLife.31657 (2018).

4 Lin, J. R., Fallahi-Sichani, M. & Sorger, P. K. Highly multiplexed imaging of single cells using a high-throughput cyclic immunofluorescence method. Nat Commun 6, 8390, doi: 10.1038/ncomms9390 (2015).

5 Dirks, R. M. & Pierce, N. A. Triggered amplification by hybridization chain reaction. Proc Natl Acad Sci U S A 101, 15275–15278, doi: 10.1073/pnas.0407024101 (2004).

6 Choi, H. M. et al. Programmable in situ amplification for multiplexed imaging of mRNA expression. Nat Biotechnol 28, 1208–1212, doi: 10.1038/nbt.1692 (2010).

7 Choi, H. M., Beck, V. A. & Pierce, N. A. Next-generation in situ hybridization chain reaction: higher gain, lower cost, greater durability. ACS Nano 8, 4284–4294, doi: 10.1021/nn405717p (2014).

8 Choi, H. M. et al. Mapping a multiplexed zoo of mRNA expression. Development 143, 3632–3637, doi: 10.1242/dev.140137 (2016).

9 Choi, J., Love, K. R., Gong, Y., Gierahn, T. M. & Love, J. C. Immuno-hybridization chain reaction for enhancing detection of individual cytokine-secreting human peripheral mononuclear cells. Anal Chem 83, 6890–6895, doi: 10.1021/ac2013916 (2011).

10 Koos, B. et al. Proximity-dependent initiation of hybridization chain reaction. Nat Commun 6, 7294, doi: 10.1038/ncomms8294 (2015).

11 Lin, R. et al. A hybridization-chain-reaction-based method for amplifying immunosignals. Nat Methods, doi: 10.1038/nmeth.4611 (2018).

12 Agasti, S. S. et al. DNA-barcoded labeling probes for highly multiplexed Exchange-PAINT imaging. Chem Sci 8, 3080–3091, doi: 10.1039/c6sc05420j (2017).

13 Callahan, L. N., Phelan, M., Mallinson, M. & Norcross, M. A. Dextran sulfate blocks antibody binding to the principal neutralizing domain of human immunodeficiency virus type 1 without interfering with gp120-CD4 interactions. J Virol 65, 1543–1550 (1991).

14 Zadeh, J. N. et al. NUPACK: Analysis and design of nucleic acid systems. J Comput Chem 32, 170–173, doi: 10.1002/jcc.21596 (2011).

15 Jungmann, R. et al. Quantitative super-resolution imaging with qPAINT. Nat Methods 13, 439–442, doi: 10.1038/nmeth.3804 (2016).

16 Coons, A. H. The demonstration of pneumococcal antigen in tissues by the use of fluorescent antibody. J Immunol 45 (1942).

17 Coons, A. H. The beginnings of immunofluorescence. J Immunol 87, 499–503 (1961).

18 Saka, S. K. et al. Immuno-SABER enables highly multiplexed and amplified protein imaging in tissues. Nat Biotechnol 37, 1080–1090, doi: 10.1038/s41587-019-0207-y (2019).

19 Goltsev, Y. et al. Deep profiling of mouse splenic architecture with CODEX multiplexed imaging. Cell 174, 968-981. e915 (2018).

20 Wang, Y. et al. Rapid Sequential in Situ Multiplexing with DNA Exchange Imaging in Neuronal Cells and Tissues. Nano Lett 17, 6131–6139, doi: 10.1021/acs.nanolett.7b02716 (2017).

21 Wang, Y. et al. Rapid sequential in situ multiplexing with DNA-Exchange-Imaging. bioRxiv, 112227 (2017).

22 Schweller, R. M. et al. Multiplexed in situ immunofluorescence using dynamic DNA complexes. Angewandte Chemie International Edition 51, 9292–9296 (2012).

23 Guo, S. M. et al. Multiplexed and high-throughput neuronal fluorescence imaging with diffusible probes. Nat Commun 10, 4377, doi: 10.1038/s41467-019-12372-6 (2019).

24 Jungmann, R. et al. Multiplexed 3D cellular super-resolution imaging with DNA-PAINT and Exchange-PAINT. Nat Methods 11, 313–318, doi: 10.1038/nmeth.2835 (2014).

25 Rosier, B. et al. Incorporation of native antibodies and Fc-fusion proteins on DNA nanostructures via a modular conjugation strategy. Chem Commun (Camb) 53, 7393–7396, doi: 10.1039/c7cc04178k (2017).

26 Popp, M. W. & Ploegh, H. L. Making and breaking peptide bonds: protein engineering using sortase. Angew Chem Int Ed Engl 50, 5024–5032, doi: 10.1002/anie.201008267 (2011).

27 Schweitzer, B. et al. Immunoassays with rolling circle DNA amplification: a versatile platform for ultrasensitive antigen detection. Proc Natl Acad Sci U S A 97, 10113–10119, doi: 10.1073/pnas.170237197 (2000).

28 Chang, J. B. et al. Iterative expansion microscopy. Nat Methods 14, 593–599, doi: 10.1038/nmeth.4261 (2017).

29 Chung, K. & Deisseroth, K. CLARITY for mapping the nervous system. Nat Methods 10, 508–513, doi: 10.1038/nmeth.2481 (2013).

30 Tillberg, P. W. et al. Protein-retention expansion microscopy of cells and tissues labeled using standard fluorescent proteins and antibodies. Nat Biotechnol 34, 987–992, doi: 10.1038/nbt.3625 (2016).

31 Chen, F. et al. Nanoscale imaging of RNA with expansion microscopy. Nat Methods 13, 679–684, doi: 10.1038/nmeth.3899 (2016).

32 Chen, F., Tillberg, P. W. & Boyden, E. S. Expansion microscopy. Science 347, 543–548, doi: 10.1126/science.1260088 (2015).

33 Stoeckius, M. et al. Simultaneous epitope and transcriptome measurement in single cells. Nat Methods 14, 865–868, doi: 10.1038/nmeth.4380 (2017).

