## Supplement figures and tables for "Multiplexed *in situ* protein imaging using DNA-barcoded antibodies with extended hybridization chain reactions"

### Supplementary Figures and Tables

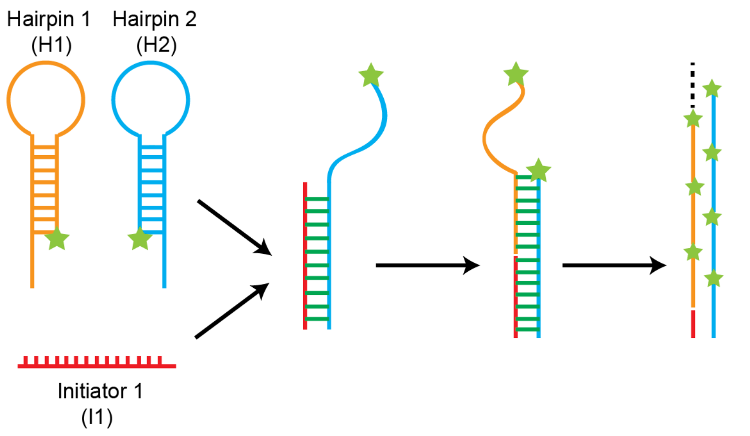


**Supplementary Figure 1. Schematic of Hybridization Chain Reaction (HCR).** The reaction includes two metastable DNA hairpin sequences (H1 and H2) and a DNA initiator sequence (I1). I1 binds to the toehold region of H2 and opens H2 via toehold displacement. The exposed sequence from H2 functions as a new initiator sequence to open hairpin H1. The process iterates to assemble a linear DNA structure. Since each hairpin is conjugated with one fluorophore, the assembled structure contains multiple fluorophores to amplify signals from a single initiator sequence.

**a** β-Tubulin (E7 antibody)- HCR-B1

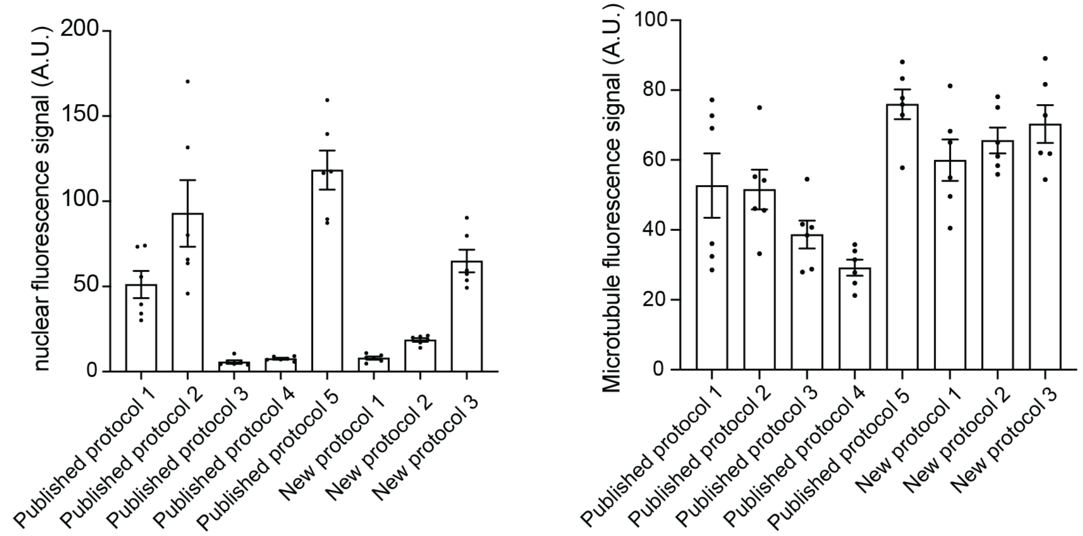


**b** α-Tubulin (YL1/2 antibody)- HCR-B1

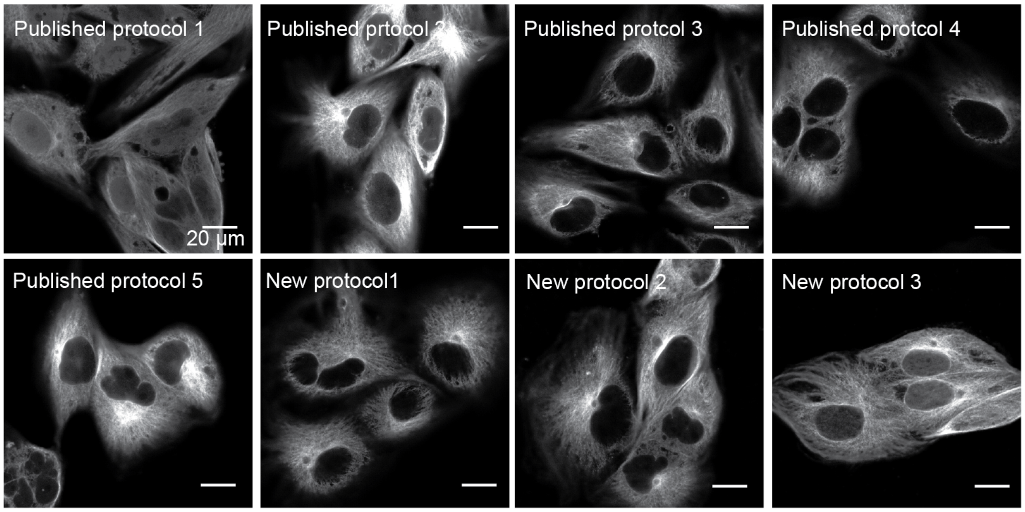


α-Tubulin (YL1/2 antibody)- HCR-B13

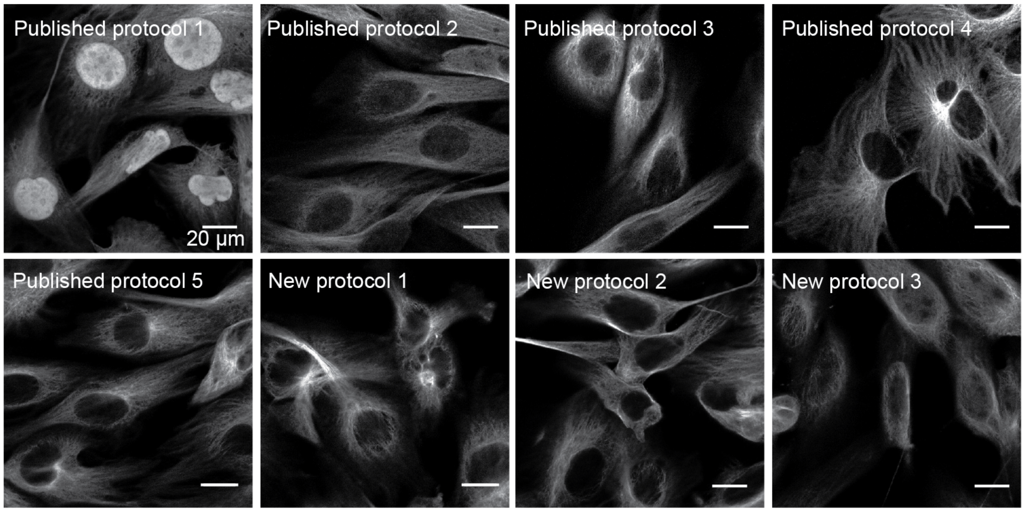

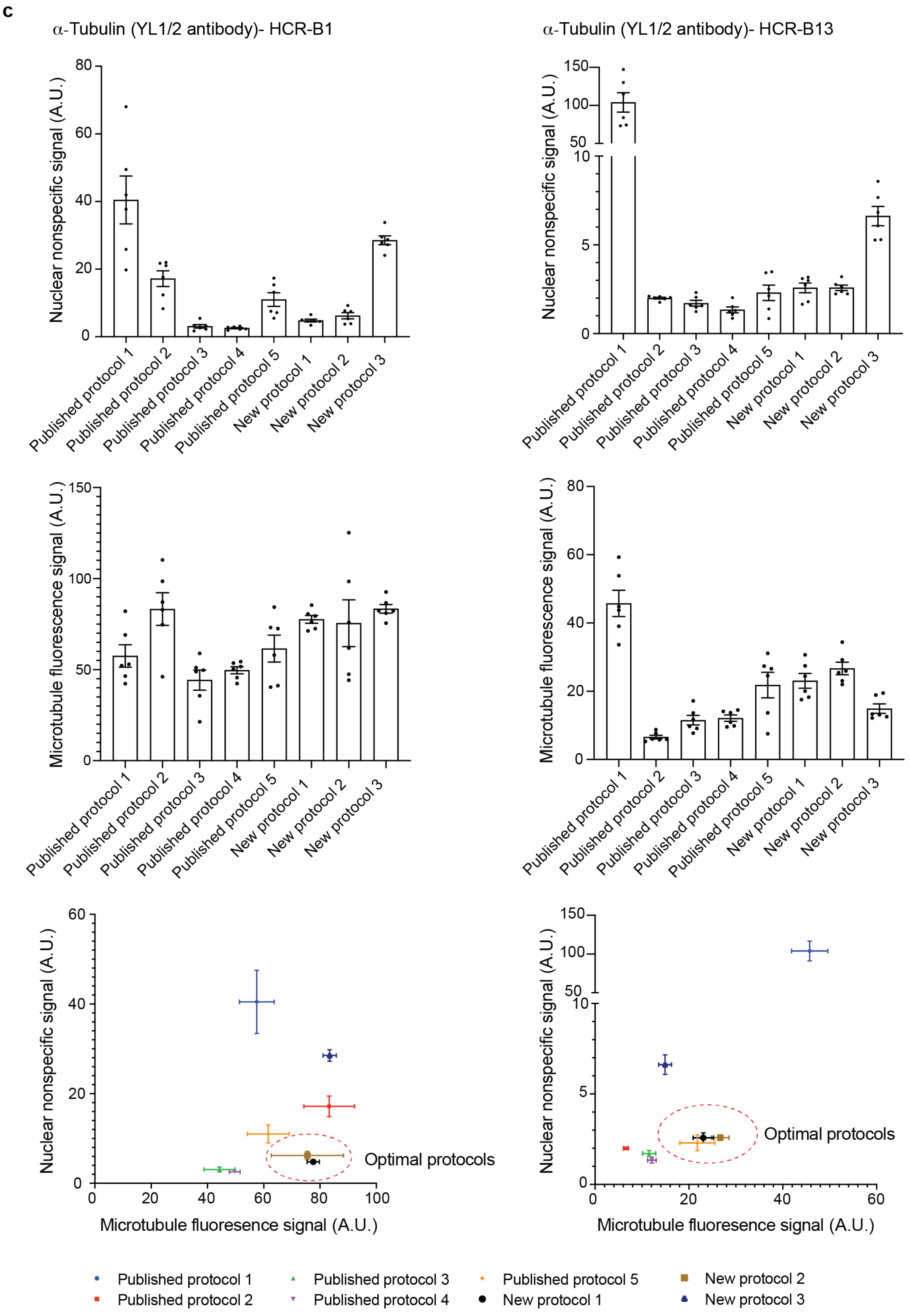
c
α-Tubulin (YL1/2 antibody)- HCR-B1
α-Tubulin (YL1/2 antibody)- HCR-B13
Nuclear nonspecific signal (A.U.)
Microtubule fluorescence signal (A.U.)
Published protocol 1
Published protocol 2
Published protocol 3
Published protocol 4
Published protocol 5
New protocol 1
New protocol 2
New protocol 3
Optimal protocols
Microtubule fluoresence signal (A.U.)
Optimal protocols
Microtubule fluoresence signal (A.U.)
Published protocol 1
Published protocol 2
Published protocol 3
Published protocol 4
Published protocol 5
New protocol 1
New protocol 2
New protocol 3

**d**

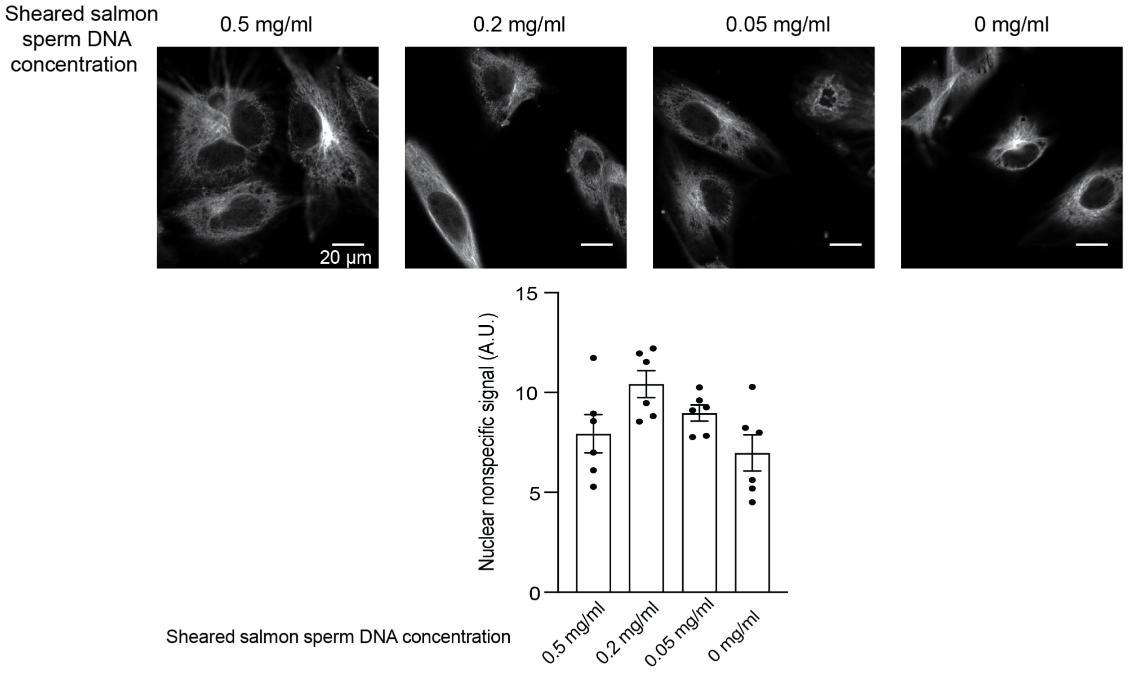

Sheared salmon sperm DNA concentration
0.5 mg/ml
0.2 mg/ml
0.05 mg/ml
0 mg/ml
20 µm
Nuclear nonspecific signal (A.U.)
15
10
5
0
Sheared salmon sperm DNA concentration
0.5 mg/ml
0.2 mg/ml
0.05 mg/ml
0 mg/ml


**e**

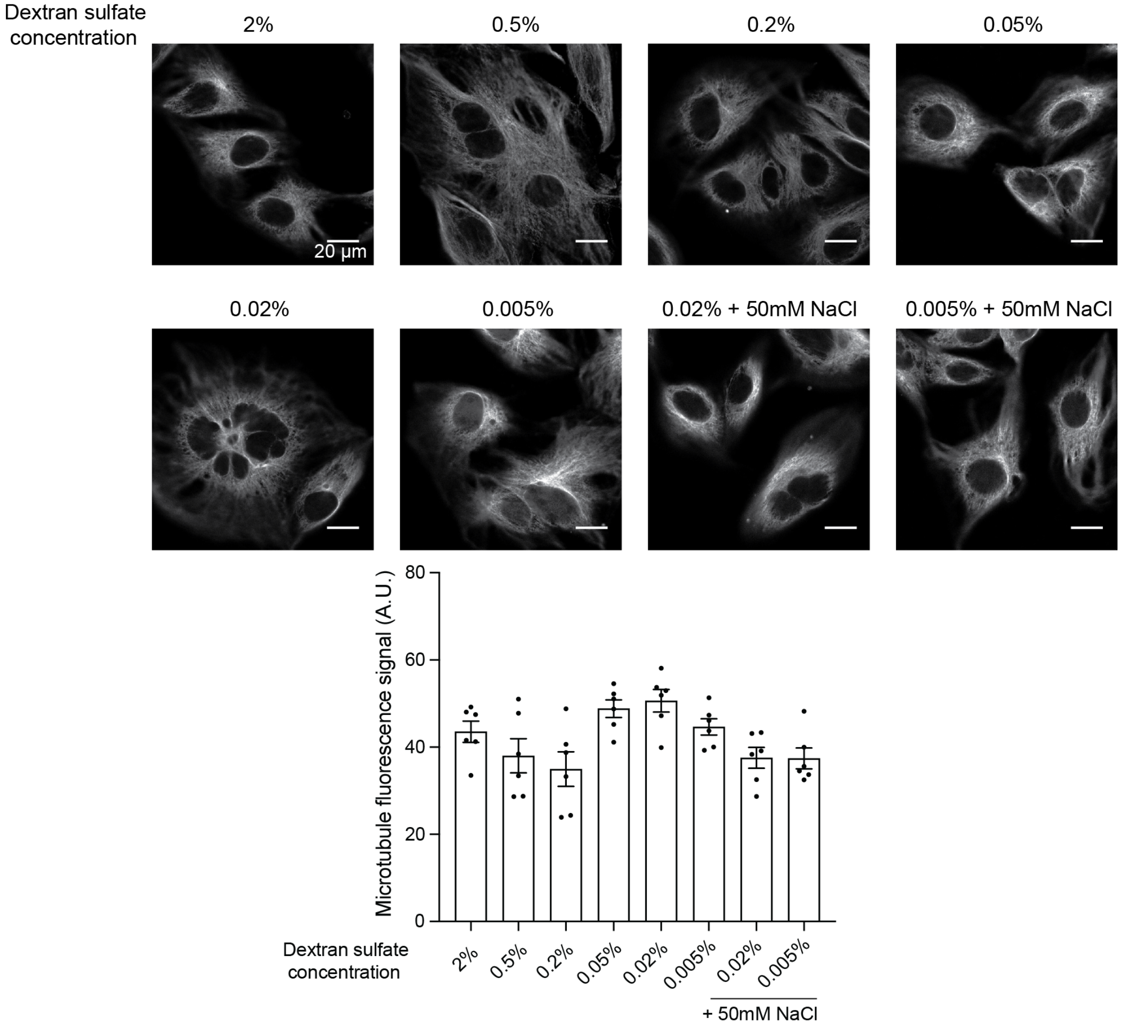

Dextran sulfate concentration
2%
0.5%
0.2%
0.05%
20 µm
0.02%
0.005%
0.02% + 50mM NaCl
0.005% + 50mM NaCl
Microtubule fluorescence signal (A.U.)
80
60
40
20
0
Dextran sulfate concentration
2%
0.5%
0.2%
0.05%
0.02%
0.005%
0.02%
0.005%
+ 50mM NaCl

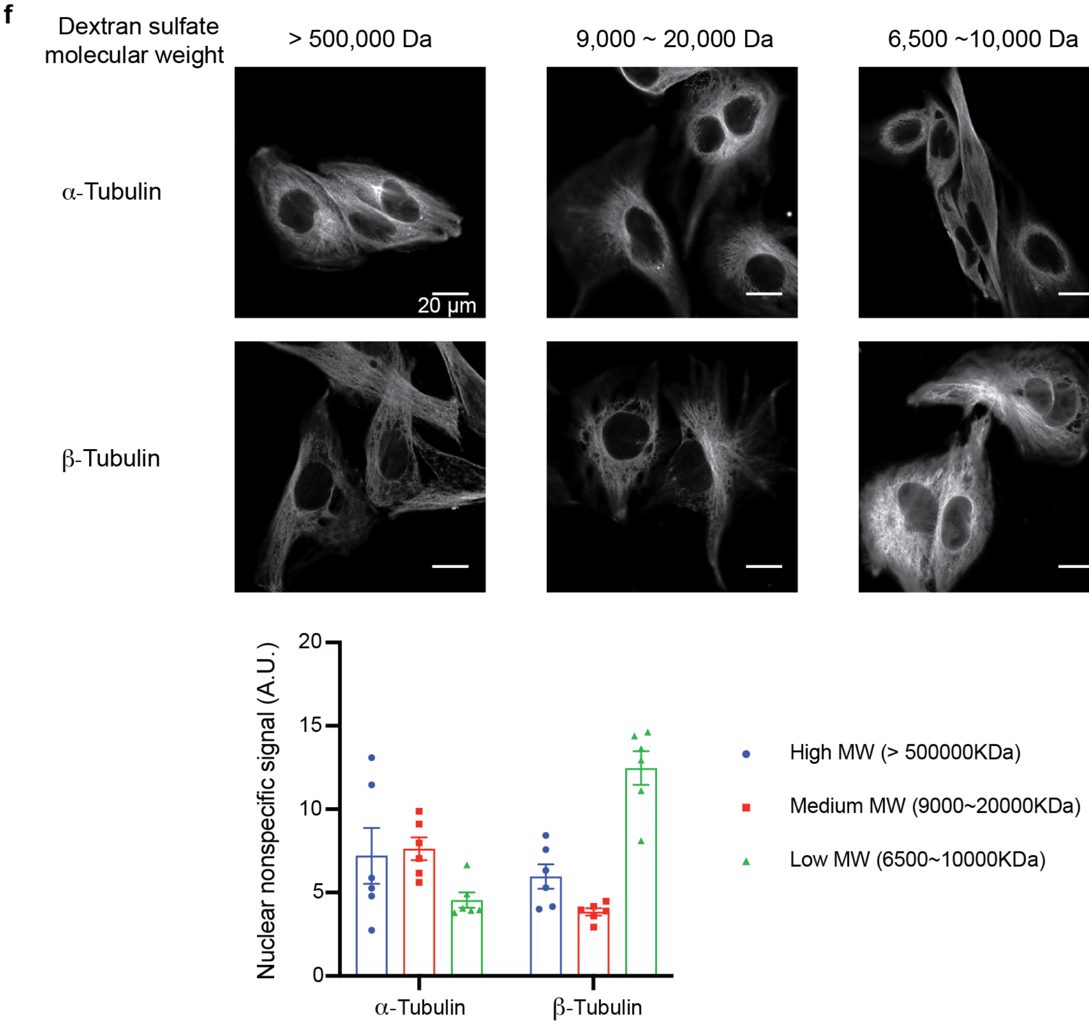


**Supplementary Figure 2**. **Development of DNA-conjugated antibody staining protocols with minimized nonspecific signals. a)** Nuclear nonspecific signals and microtubule fluorescence signals used to plot the 2-D plot in Figure 2d. Error bar is SEM and n = 6 areas. **b)** Comparison of different antibody staining protocols for nonspecific nuclear signal reduction efficiency. BS-C-1 cells were stained with HCR-B1I1 or B13I1-conjugated anti-α-Tubulin YL1/2 antibodies with protocols described in Supplementary Table 1. **c)** Quantification of nuclear nonspecific signals and microtubule signals in panel b. Error bar is SEM and n = 6 areas. **d)** Titration of sheared salmon sperm DNA in the blocking and antibody incubation buffer, and quantification of nuclear nonspecific signals. BS-C-1 cells were stained with HCR-B1I1-conjugated anti-β-Tubulin E7 antibodies. Error bar is SEM and n = 6 areas. **e)** Titration of dextran sulfate in the antibody incubation buffer, and quantification of microtubule fluorescence signals. The nuclear nonspecific signals were also quantified and shown in Figure 2e. BS-C-1 cells were stained with HCR-B1I1-conjugated anti-β-Tubulin E7 antibodies. Error bar is SEM and n = 6 areas. f) Comparison of dextran sulfate with different molecular weight for nonspecific nuclear signal reduction efficiency. BS-C-1 cells were stained with HCR-B1I1-conjugated anti-α-Tubulin YL1/2 or anti-β-Tubulin E7 antibodies. Error bar is SEM and n = 6 areas.

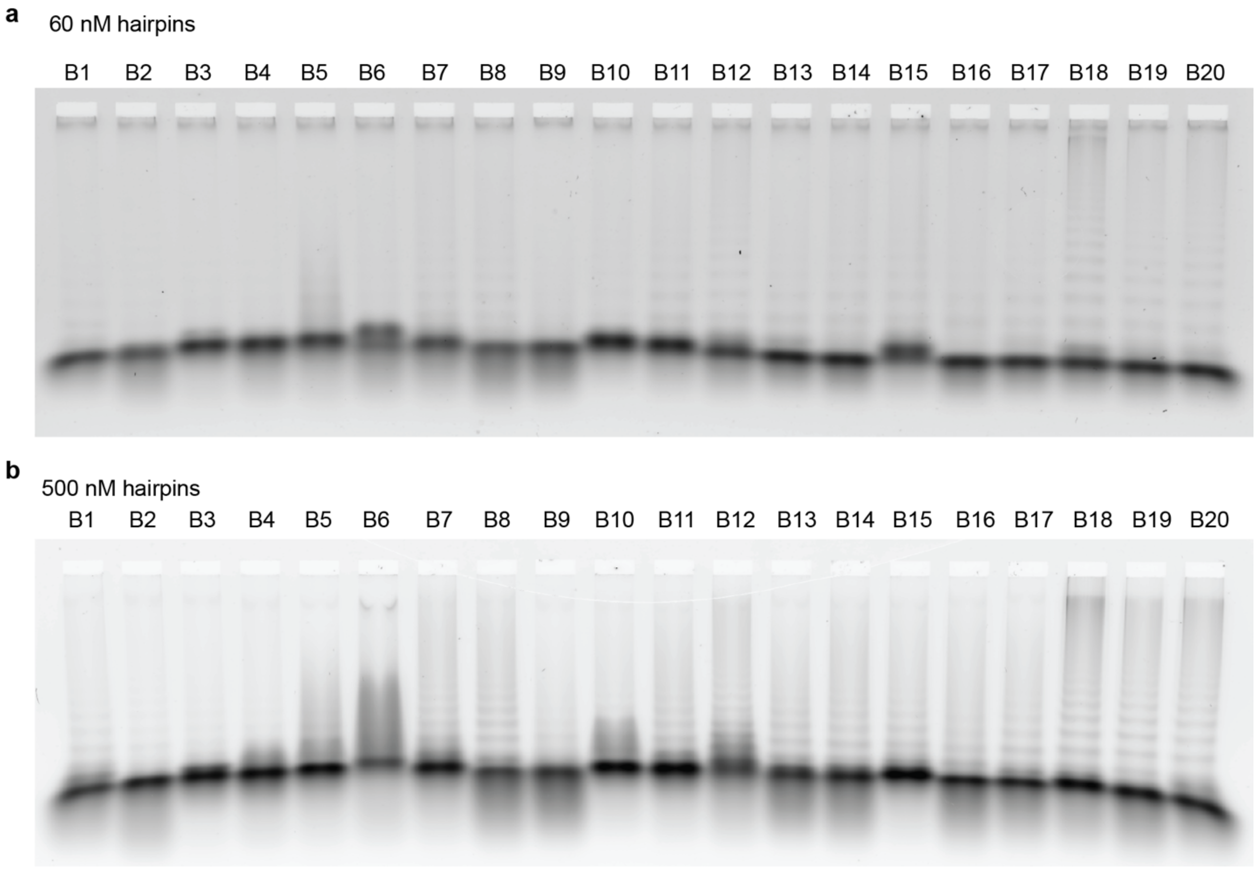


**Supplementary Figure 3. *In vitro* leakage analysis of newly designed additional HCR pairs.** For each pair, either 60 nM (a) or 500 nM (b) hairpins (H1 + H2) were added in a PCR tube without HCR initiator and left at room temperature for 24 hours. The leakage results were visualized by running the products on agarose gels.

**HCR hairpins crosstalk analysis**:

**B1 hairpins (B1H1+B1H2) with indicated initiators** (B2 left lane is B2I1; B2 right lane is B2I2; \ is negative control without initiator, last lane is positive control with B1I1; B2-B11 are in gel 1 and B12-B1 are in gel 2):

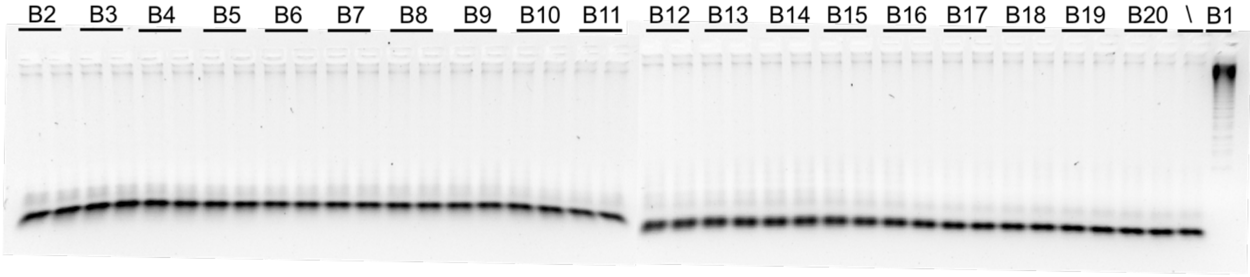


**B2 hairpins (B2H1+B2H2) with indicated initiators** (B1 left lane is B1I1; B1 right lane is B1I2; \ is negative control without initiator, last lane is positive control with B2I1; B1-B11 are in gel 1 and B12-B2 are in gel 2):

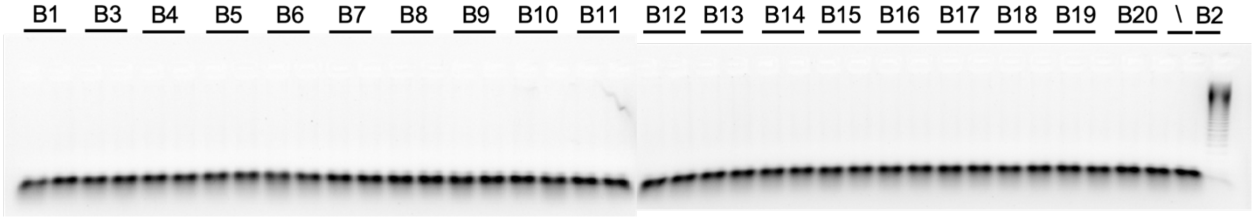


**B3 hairpins (B3H1+B3H2) with indicated initiators** (B1 left lane is B1I1; B1 right lane is B1I2; \ is negative control without initiator, last lane is positive control with B3I1; B1-B11 are in gel 1 and B12-B3 are in gel 2):

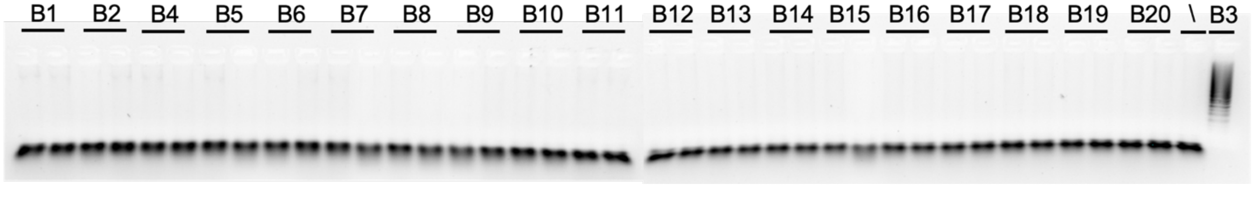

**B4 hairpins (B4H1+B4H2) with indicated initiators** (B1 left lane is B1I1; B1 right lane is B1I2; \ is negative control without initiator, last lane is positive control with B4I1; B1-B11 are in gel 1 and B12-B4 are in gel 2):

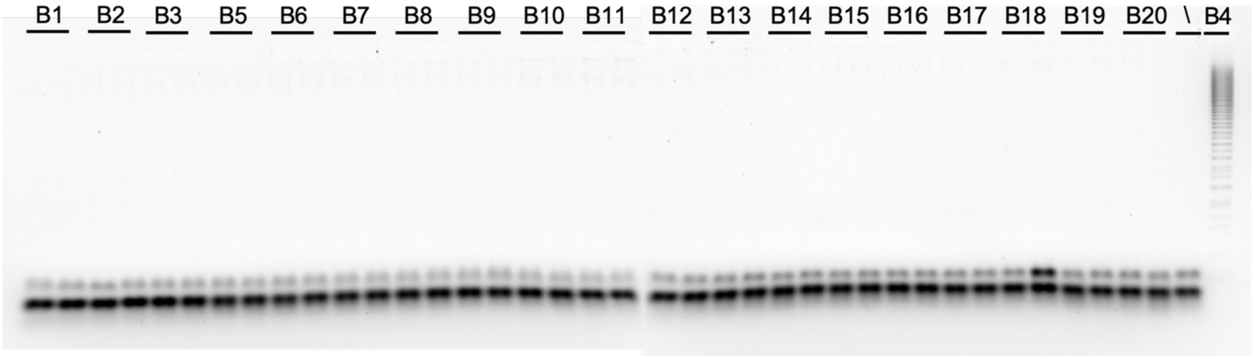


**B5 hairpins (B5H1+B5H2) with indicated initiators** (B1 left lane is B1I1; B1 right lane is B1I2; \ is negative control without initiator, last lane is positive control with B5I1; B1-B11 are in gel 1 and B12-B5 are in gel 2):

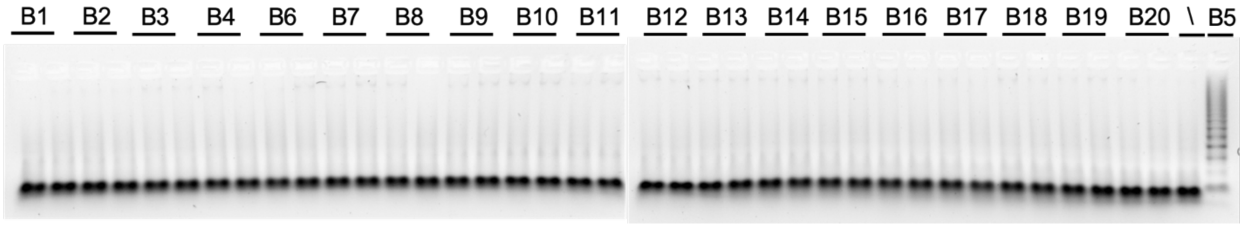


**B7 hairpins (B7H1+B7H2) with indicated initiators** (B1 left lane is B1I1; B1 right lane is B1I2; \ is negative control without initiator, last lane is positive control with B7I1; crosstalk lane was marked with a red box; B1-B11 are in gel 1 and B12-B7 are in gel 2):

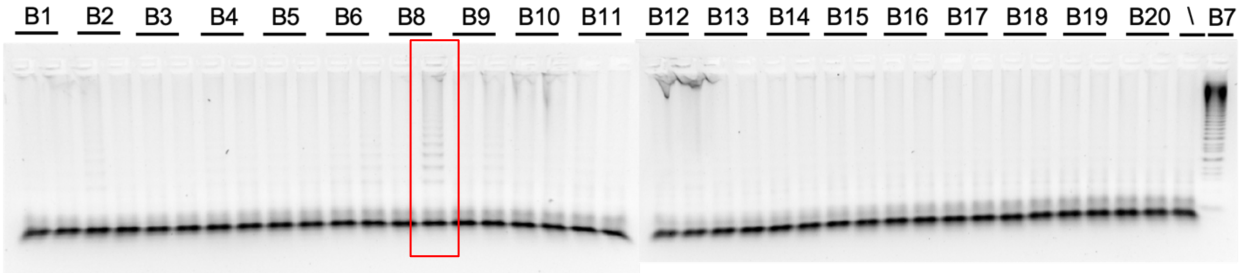

**B8 hairpins (B8H1+B8H2) with indicated initiators** (B1 left lane is B1I1; B1 right lane is B1I2; \ is negative control without initiator, last lane is positive control with B8I1; B1-B11 are in gel 1 and B12-B8 are in gel 2):

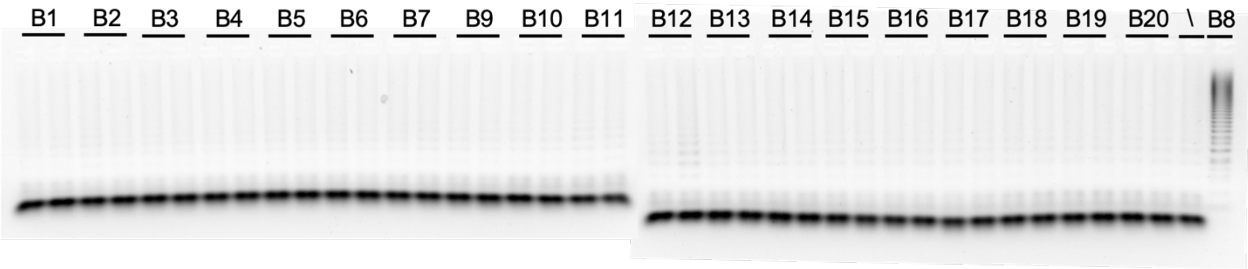


**B9 hairpins (B9H1+B9H2) with indicated initiators** (B1 left lane is B1I1; B1 right lane is B1I2; \ is negative control without initiator, last lane is positive control with B9I1; B1-B11 are in gel 1 and B12-B9 are in gel 2):

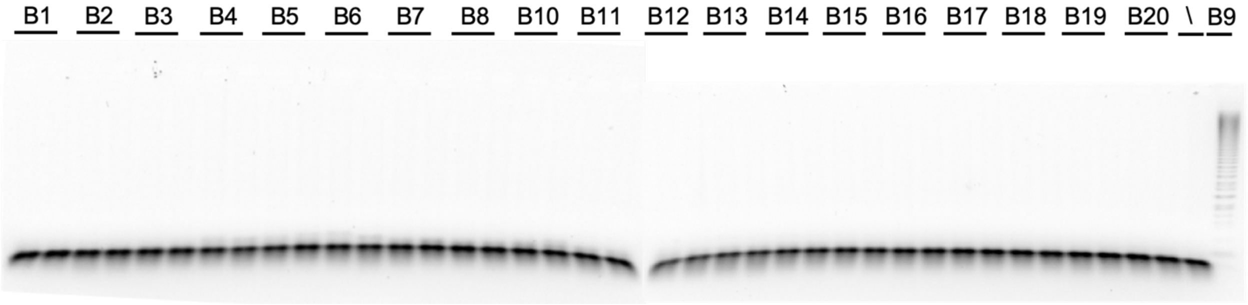


**B10 hairpins (B10H1+B10H2) with indicated initiators** (B1 left lane is B1I1; B1 right lane is B1I2; \ is negative control without initiator, last lane is positive control with B10I1; B1-B11 are in gel 1 and B12-B10 are in gel 2):

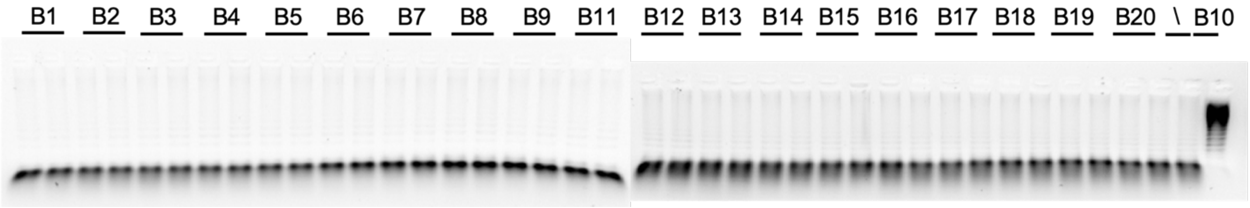

**B11 hairpins (B11H1+B11H2) with indicated initiators** (B1 left lane is B1I1; B1 right lane is B1I2; \ is negative control without initiator, last lane is positive control with B11I1; B1-B10 are in gel 1 and B12-B11 are in gel 2):

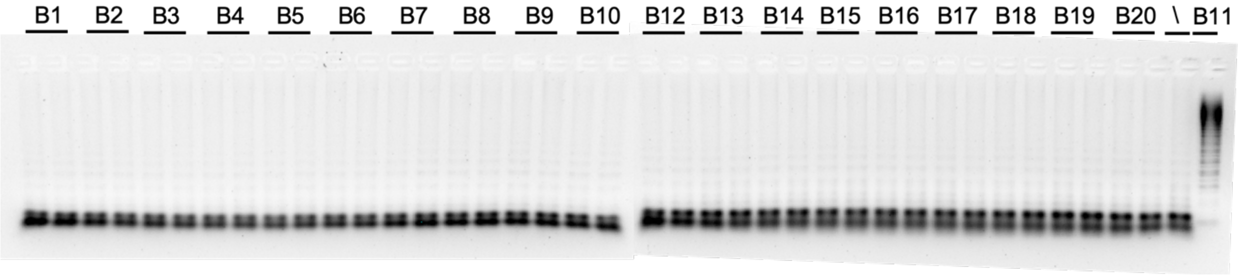


**B13 hairpins (B13H1+B13H2) with indicated initiators** (B1 left lane is B1I1; B1 right lane is B1I2; \ is negative control without initiator, last lane is positive control with B13I1; B1-B10 are in gel 1 and B11-B13 are in gel 2):

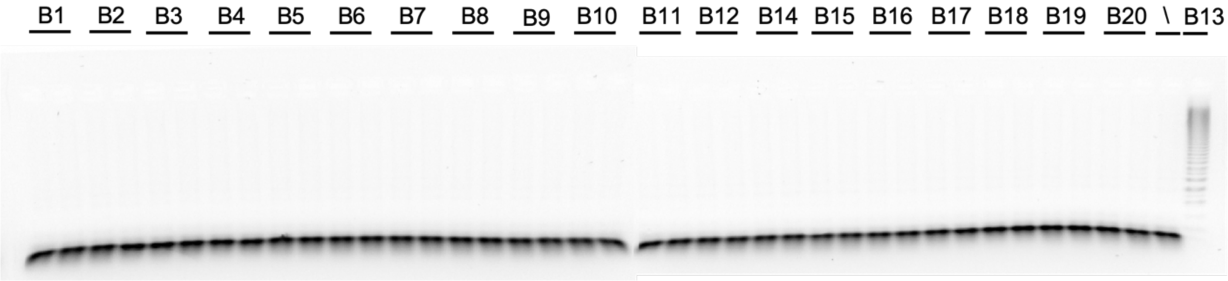


**B14 hairpins (B14H1+B14H2) with indicated initiators** (B1 left lane is B1I1; B1 right lane is B1I2; \ is negative control without initiator, last lane is positive control with B14I1; B1-B10 are in gel 1 and B11-B14 are in gel 2):

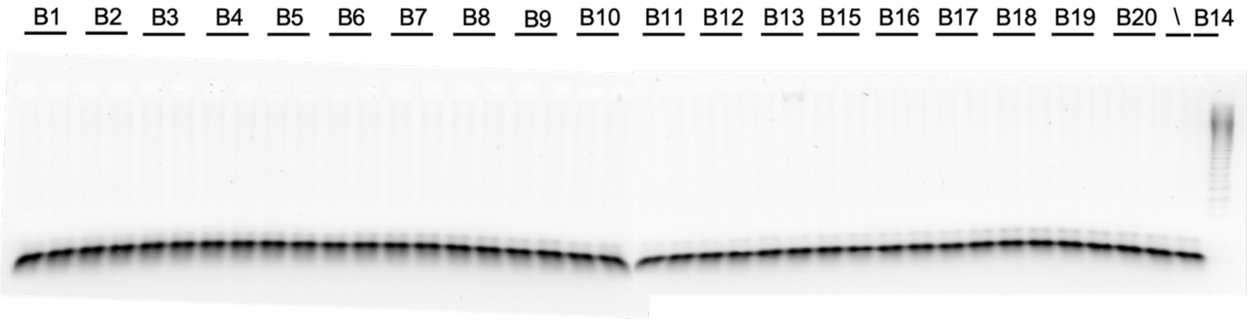

**B15 hairpins (B15H1+B15H2) with indicated initiators** (B1 left lane is B1I1; B1 right lane is B1I2; \ is negative control without initiator, last lane is positive control with B15I1; B1-B10 are in gel 1 and B11-B15 are in gel 2):

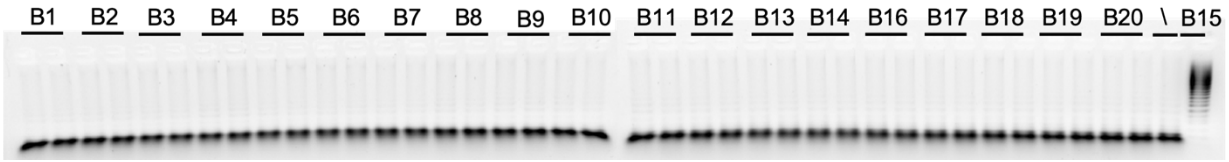


**B16 hairpins (B16H1+B16H2) with indicated initiators** (B1 left lane is B1I1; B1 right lane is B1I2; \ is negative control without initiator, last lane is positive control with B16I1; crosstalk lane was marked with a red box; B1-B10 are in gel 1 and B11-B16 are in gel 2):

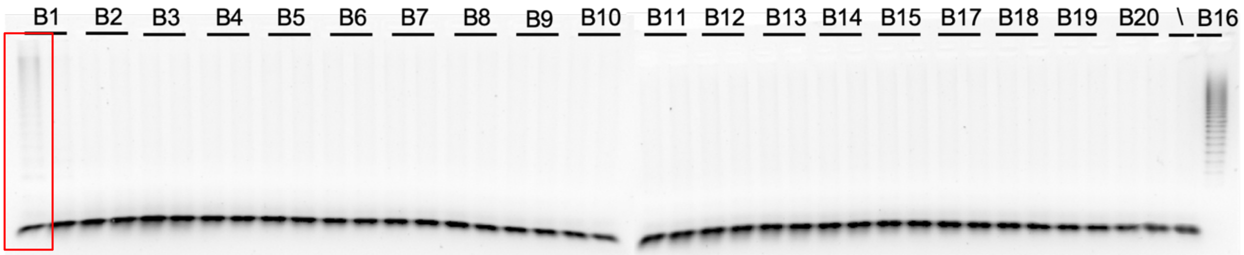


**B17 hairpins (B17H1+B17H2) with indicated initiators** (B1 left lane is B1I1; B1 right lane is B1I2; \ is negative control without initiator, last lane is positive control with B17I1; B1-B10 are in gel 1 and B11-B17 are in gel 2):

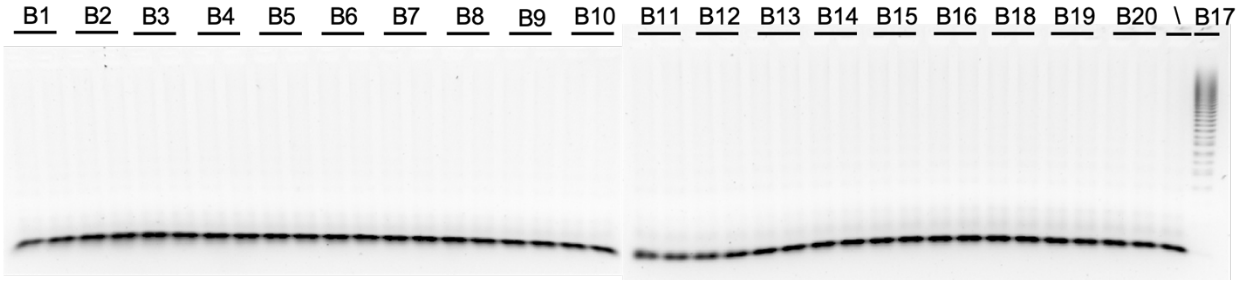


**Supplementary Figure 4. *In vitro* pairwise crosstalk analysis of 15 HCR pairs (excluding B6, B12, B18-20).** For each reaction, 500 nM of each of the HCR hairpin pairs (H1 + H2) was added in a PCR tube with 50 nM indicated initiator sequence, and left at room temperature for 24 hours to react. The products were visualized on agarose gels. The \ lane in each gel was a negative control in which no initiator was added, whereas the last lane in each gel was a positive control in which the cognate initiator was added. Crosstalk lanes were marked with red boxes.

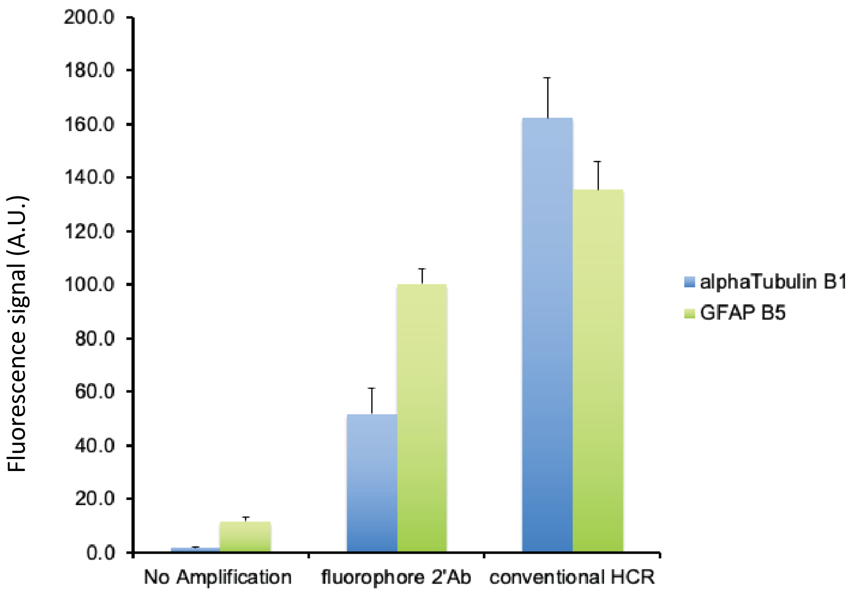


**Supplementary Figure 5. Quantification of image fluorescence intensities of α-Tubulin and GFAP from Figure 4a.** Error bar is SEM and n = 12 areas.

**Supplementary Table 1. Buffer composition for different blocking and antibody incubation buffers used in Figure 2b.** The compositions are derived from the published protocols listed below or are newly designed.

| | **Blocking buffer** (in 1 × PBS) | **Antibody incubation buffer** (in 1 × PBS) |
|---|---|---|
| Published protocol 1 (Lin et al. 2018) | 5% BSA + 0.1% Triton | 3% BSA + 0.1% Triton |
| Published protocol 2 (Guo et al. 2019) | 5% BSA + 0.1% Triton + 1 mg/ml sheared sperm DNA | 3% BSA + 0.1% Triton + 1 mg/ml sheared sperm DNA |
| Published protocol 3 (Lee et al. 2018) | 5% BSA + 0.1% Triton + 1 mg/ml sheared sperm DNA + 0.5% dextran sulfate | 5% BSA + 0.1% Triton + 1 mg/ml sheared sperm DNA + 0.5% dextran sulfate |
| Published protocol 4 (Chen et al. 2015) | 5% normal serum | 5% normal serum + 0.1% Triton + 1 mg/ml sheared sperm DNA + 10% dextran sulfate in 2× SSC |
| Published protocol 5 (Schürch et al. 2019) | 10 µM complementary DNA sequence + 0.1 mg/ml IgG + 0.5 mg/ml sheared sperm DNA + 0.1% Triton | 10 µM complementary DNA sequence + 0.1 mg/ml IgG + 0.5 mg/ml sheared sperm DNA + 0.1% Triton |
| New protocol 1 | 10 µM complementary DNA sequence + 5% BSA + 0.5 mg/ml sheared sperm DNA + 0.1% Triton | 10 µM complementary DNA sequence + 5% BSA + 0.5 mg/ml sheared sperm DNA + 0.1% Triton + 0.5% dextran sulfate |
| New protocol 2 | 5% BSA + 0.1% Triton | 3% BSA + 0.1% Triton + 10 µM complementary DNA sequence + 0.5% dextran sulfate |
| New protocol 3 | 5% BSA + 0.1% Triton | 3% BSA + 0.1% Triton + 10 µM complementary DNA sequence + 250 mM NaCl |

*Published protocol 1:*
Lin, R., Feng, Q., Li, P., … and Luo, M. (2018). A hybridization-chain-reaction-based method for amplifying immunosignals. *Nature methods*, *15*(4), p.275.

*Published protocol 2:*
Guo, S.M., Veneziano, R., Gordonov, S., … and Bathe, M. (2019). Multiplexed and high-throughput neuronal fluorescence imaging with diffusible probes. *Nature communications*, 10(1), pp.1-14.

*Published protocol 3:*
Lee, J., Geiss, G.K., Demirkan, G., … and Mills, G.B. (2018). Implementation of a multiplex and quantitative proteomics platform for assessing protein lysates using DNA-barcoded antibodies. *Molecular & Cellular Proteomics*, *17*(6), pp.1245-1258.

*Published protocol 4:*
Chen, F., Tillberg, P.W. and Boyden, E.S. (2015). Expansion microscopy. *Science*, *347*(6221), pp.543-548.

*Published protocol 5:*
Schürch, C.M., Bhate, S., Barlow, G.L., … and Nolan Garry (2019). Coordinated cellular neighborhoods orchestrate antitumoral immunity at the colorectal cancer invasive front. *CELL-D-19-02119*.

**Supplementary Table 2. DNA sequences of hairpins and initiators for all 20 HCR pairs.**

| Name | DNA sequence |
|---|---|
| B1H1 | CGTAAAGGAAGACTCTTCCCGTTTGCTGCCCTCCTCGCATTCTTTCTTGAGGAGGGCAGCAAACGGGAAGAG |
| B1H2 | GAGGAGGGCAGCAAACGGGAAGAGTCTTCCTTTACGCTCTTCCCGTTTGCTGCCCTCCTCAAGAAAGAATGC |
| B1I1 | GCATTCTTTCTTGAGGAGGGCAGCAAACGGGAAGAG |
| B1I2 | GAGGAGGGCAGCAAACGGGAAGAGTCTTCCTTTACG |
| B2H1 | CCTCGTAAATCCTCATCAATCATCCAGTAAACCGCCGATGATTGATGAGGATTTACGAGGATGGACTGAGCT |
| B2H2 | GGCGGTTTACTGGATGATTGATGAGGATTTACGAGGAGCTCAGTCCATCCTCGTAAATCCTCATCAATCATC |
| B2I1 | AGCTCAGTCCATCCTCGTAAATCCTCATCAATCATC |
| B2I2 | CCTCGTAAATCCTCATCAATCATCCAGTAAACCGCC |
| B3H1 | GTCCCTGCCTCTATATCTCCACTCAACTTTAACCCGGAGTGGAGATATAGAGGCAGGGACGGATTAGACTTT |
| B3H2 | CGGGTTAAAGTTGAGTGGAGATATAGAGGCAGGGACAAAGTCTAATCCGTCCCTGCCTCTATATCTCCACTC |
| B3I1 | AAAGTCTAATCCGTCCCTGCCTCTATATCTCCACTC |
| B3I2 | GTCCCTGCCTCTATATCTCCACTCAACTTTAACCCG |
| B4H1 | CCTCAACCTACCTCCAACTCTCACCATATTCGCTTCGTGAGAGTTGGAGGTAGGTTGAGGTCTGTAAATGTG |
| B4H2 | GAAGCGAATATGGTGAGAGTTGGAGGTAGGTTGAGGCACATTTACAGACCTCAACCTACCTCCAACTCTCAC |
| B4I1 | CACATTTACAGACCTCAACCTACCTCCAACTCTCAC |
| B4I2 | CCTCAACCTACCTCCAACTCTCACCATATTCGCTTC |
| B5H1 | ATTGGATTTGTAGGGTAGATAGAGATTGGGAGTGAGCACTTCATATCACTCACTCCCAATCTCTATCTACCC |
| B5H2 | CTCACTCCCAATCTCTATCTACCCTACAAATCCAATGGGTAGATAGAGATTGGGAGTGAGTGATATGAAGTG |
| B5I1 | CACTTCATATCACTCACTCCCAATCTCTATCTACCC |
| B5I2 | CTCACTCCCAATCTCTATCTACCCTACAAATCCAAT |
| B6H1 | CACCCTCACTACCTCACCCTCACCTCCATCCACCTCGGTGAGGGTGAGGTAGTGAGGGTGAGTGGGAATGGG |
| B6H2 | GAGGTGGATGGAGGTGAGGGTGAGGTAGTGAGGGTGCCCATTCCCACTCACCCTCACTACCTCACCCTCACC |
| B6I1 | CCCATTCCCACTCACCCTCACTACCTCACCCTCACC |
| B6I2 | CACCCTCACTACCTCACCCTCACCTCCATCCACCTC |
| B7H1 | CCTACCTCCAATCCCTACCCTCACTACATCCTCACCGTGAGGGTAGGGATTGGAGGTAGGTGGAGGTTGAAG |
| B7H2 | GGTGAGGATGTAGTGAGGGTAGGGATTGGAGGTAGGCTTCAACCTCCACCTACCTCCAATCCCTACCCTCAC |
| B7I1 | CTTCAACCTCCACCTACCTCCAATCCCTACCCTCAC |
| B7I2 | CCTACCTCCAATCCCTACCCTCACTACATCCTCACC |
| B8H1 | CGTCTCCATCACTCGCACTCTACCGTCAAGTCAAACGGTAGAGTGCGAGTGATGGAGACGAGATAATCAAGG |
| B8H2 | GTTTGACTTGACGGTAGAGTGCGAGTGATGGAGACGCCTTGATTATCTCGTCTCCATCACTCGCACTCTACC |
| B8I1 | CCTTGATTATCTCGTCTCCATCACTCGCACTCTACC |
| B8I2 | CGTCTCCATCACTCGCACTCTACCGTCAAGTCAAAC |

| | |
|---|---|
| B9H1 | CCACTCTCAGCACACTCCCAACCCTACTACAAGCTCGGGTTGGGAGTGTGCTGAGA GTGGAGTAGATACGTG |
| B9H2 | GAGCTTGTAGTAGGGTTGGGAGTGTGCTGAGAGTGGCACGTATCTACTCCACTCTC AGCACACTCCCAACCC |
| B9I1 | CACGTATCTACTCCACTCTCAGCACACTCCCAACCC |
| B9I2 | CCACTCTCAGCACACTCCCAACCCTACTACAAGCTC |
| B10H1 | CCTCTACCTACTCGACTACCCTAGCCGTAACTTCACCTAGGGTAGTCGAGTAGGTA GAGGAGTATCTTGAGG |
| B10H2 | GTGAAGTTACGGCTAGGGTAGTCGAGTAGGTAGAGGCCTCAAGATACTCCTCTACC TACTCGACTACCCTAG |
| B10I1 | CCTCAAGATACTCCTCTACCTACTCGACTACCCTAG |
| B10I2 | CCTCTACCTACTCGACTACCCTAGCCGTAACTTCAC |
| B11H1 | ACTCCTACGTCGACCACACTCATCCTGCATGTTCCCGATGAGTGTGGTCGACGTAG GAGTGATATCTAAGCG |
| B11H2 | GGGAACATGCAGGATGAGTGTGGTCGACGTAGGAGTCGCTTAGATATCACTCCTA CGTCGACCACACTCATC |
| B11I1 | CGCTTAGATATCACTCCTACGTCGACCACACTCATC |
| B11I2 | ACTCCTACGTCGACCACACTCATCCTGCATGTTCCC |
| B12H1 | GCTCTCATGCCTCTCGTACTACCGTCACAAGTAAACCGGTAGTACGAGAGGCATGA GAGCGATAGCGTAAGG |
| B12H2 | GTTTACTTGTGACGGTAGTACGAGAGGCATGAGAGCCCTTACGCTATCGCTCTCAT GCCTCTCGTACTACCG |
| B12I1 | CCTTACGCTATCGCTCTCATGCCTCTCGTACTACCG |
| B12I2 | GCTCTCATGCCTCTCGTACTACCGTCACAAGTAAAC |
| B13H1 | CCTGCTTTATGCTCAACATACAACCAGAAATGCGGCGTTGTATGTTGAGCATAAAG CAGGAAGGCGTTACCT |
| B13H2 | GCCGCATTTCTGGTTGTATGTTGAGCATAAAGCAGGAGGTAACGCCTTCCTGCTTT ATGCTCAACATACAAC |
| B13I1 | AGGTAACGCCTTCCTGCTTTATGCTCAACATACAAC |
| B13I2 | CCTGCTTTATGCTCAACATACAACCAGAAATGCGGC |
| B14H1 | GAGCGACCCTATATTTCTGCACAGAAGTTATACCGGCTGTGCAGAAATATAGGGTC GCTCGCTATTGACATT |
| B14H2 | CCGGTATAACTTCTGTGCAGAAATATAGGGTCGCTCAATGTCAATAGCGAGCGACC CTATATTTCTGCACAG |
| B14I1 | AATGTCAATAGCGAGCGACCCTATATTTCTGCACAG |
| B14I2 | GAGCGACCCTATATTTCTGCACAGAAGTTATACCGG |
| B15H1 | CCACAAGGTATCTCGAACACTCTCCAAATTGGCTACGAGAGTGTTCGAGATACCTT GTGGTGTGTTAATCTG |
| B15H2 | GTAGCCAATTTGGAGAGTGTTCGAGATACCTTGTGGCAGATTAACACACCACAAG GTATCTCGAACACTCTC |
| B15I1 | CAGATTAACACACCACAAGGTATCTCGAACACTCTC |
| B15I2 | CCACAAGGTATCTCGAACACTCTCCAAATTGGCTAC |
| B16H1 | GACGAGCGCACCAATCGGCAAGTGTCATGGATTAGGCACTTGCCGATTGGTGCGCT CGTCATGAATGAAAGC |
| B16H2 | CCTAATCCATGACACTTGCCGATTGGTGCGCTCGTCGCTTTCATTCATGACGAGCG CACCAATCGGCAAGTG |
| B16I1 | GCTTTCATTCATGACGAGCGCACCAATCGGCAAGTG |
| B16I2 | GACGAGCGCACCAATCGGCAAGTGTCATGGATTAGG |
| B17H1 | GTGGACACCTGCTAATCGGATGAGTGTTCGTTATCGCTCATCCGATTAGCAGGTGT CCACAACAAACAATCG |

| | |
|---|---|
| B17H2 | CGATAACGAACACTCATCCGATTAGCAGGTGTCCACCGATTGTTTGTTGTGGACAC CTGCTAATCGGATGAG |
| B17I1 | CGATTGTTTGTTGTGGACGCATGCTAATCGGATGAG |
| B17I2 | GTGGACGCATGCTAATCGGATGAGTGTTCGTTATCG |
| B18H1 | GGACGATAAACCTGATCTATCTTCCTGTATAGGCTCGAAGATAGATCAGGTTTATC GTCCATCGTCACTGGT |
| B18H2 | GAGCCTATACAGGAAGATAGATCAGGTTTATCGTCCACCAGTGACGATGGACGAT AAACCTGATCTATCTTC |
| B18I1 | ACCAGTGACGATGGACGATAAACCTGATCTATCTTC |
| B18I2 | GGACGATAAACCTGATCTATCTTCCTGTATAGGCTC |
| B19H1 | GACCTACGGACTTAATCTGCTGTCATCTTAAAGCGGGACAGCAGATTAAGTCCGTA GGTCCGTTCACACTAT |
| B19H2 | CCGCTTTAAGATGACAGCAGATTAAGTCCGTAGGTCATAGTGTGAACGGACCTACG GACTTAATCTGCTGTC |
| B19I1 | ATAGTGTGAACGGACCTACGGACTTAATCTGCTGTC |
| B19I2 | GACCTACGGACTTAATCTGCTGTCATCTTAAAGCGG |
| B20H1 | TCAGTACCTTGCTCGTTCTAACATCTATTTGTCTACATGTTAGAACGAGCAAGGTAC TGATGAGTAATACTG |
| B20H2 | GTAGACAAATAGATGTTAGAACGAGCAAGGTACTGACAGTATTACTCATCAGTAC CTTGCTCGTTCTAACAT |
| B20I1 | CAGTATTACTCATCAGTACCTTGCTCGTTCTAACAT |
| B20I2 | TCAGTACCTTGCTCGTTCTAACATCTATTTGTCTAC |
